# Single-cell transcriptomics reveals probiotic reversal of neonatal morphine-induced gene disruptions underlying adolescent pain hypersensitivity

**DOI:** 10.1101/2025.05.30.657034

**Authors:** Junyi Tao, Danielle Antoine, Richa Jalodia, Eridania Valdes, Sean Michael Boyles, William Hulme, Sabita Roy

**Author notes:** Authors Contributed Equally (Junyi Tao, Danielle Antoine). Corresponding author Sabita Roy, Ph.D. 1501 NW 10^th^ Ave, BRB 531 Miami, FL 33136.

## Abstract

Neonatal morphine is commonly administered in the Neonatal Intensive Care Unit (NICU) to manage pain. However, its long-term effects on neurodevelopment of pain pathways, remain a significant concern. The midbrain is a core region that plays a central role in pain processing and opioid-mediated analgesia. Here, we performed single-cell RNA sequencing to study gene expression in 107,427 midbrain single cells from adolescent mice neonatally exposed to either saline, morphine, or morphine with the probiotic *Bifidobacterium infantis* (*B. infantis*). We found broad alterations in transcriptomics within neurons, astrocytes, oligodendrocytes, and microglial cells. Analysis of differentially regulated genes revealed down regulation of HOX genes and upregulation of pathways related to neurotransmitter signaling and pain in adolescence that were neonatally treated with morphine. Interestingly, neonatal probiotic supplementation mitigated these morphine-induced alterations on the transcriptome. This study presents the first single-cell RNA sequencing dataset of the adolescent midbrain following neonatal morphine exposure and probiotic intervention. These findings offer new insights into the neurodevelopmental impact of early opioid exposure and highlight the therapeutic potential of microbiome-targeted interventions.

## Introduction

Opioid analgesics, such as morphine, are frequently administered in Neonatal Intensive Care Units (NICUs) to manage moderate to severe pain in neonates undergoing surgery or invasive procedures^1, 2^. Despite their widespread use, the long-term neurobiological consequences of neonatal morphine exposure (NME) remain a critical area of investigation due to the potential for adverse outcomes, including an increased risk of developing opioid use disorders. Emerging clinical and preclinical evidence indicates that NME may lead to detrimental neurodevelopmental effects, such as altered pain sensitivity, changes in locomotor activity, and impairments in cognition and learning^2, 3^.

We previously demonstrated that NME induces persistent thermal and mechanical pain hypersensitivity in adolescent mice, , driven by gut microbial dysbiosis, intestinal barrier dysfunction, and systemic inflammation^4^. Notably, neonatal supplementation with the probiotic *Bifidobacterium infantis* (*B. infantis*) prevented the development of pain hypersensitivity by restoring gut microbial dysbiosis and the associated inflammation. Additonally, bulk RNA sequencing of midbrain tissue revealed upregulation of gene networks related to excitatory signaling and neuroimmune activation, many of which were normalized by probiotic treatment which implicated the gut-brain axis as a key driver of long-term alterations in pain processing following early-life opioid exposure^4^.

Despite previous findings, the cellular specificity and mechanistic basis of transcriptional changes in the brain remain poorly understood. The midbrain, which encompasses key nodes such as the periaqueductal gray (PAG) and ventral tegmental area (VTA), plays a central role in pain modulation and reward circuitry^5, 6^. However, bulk tissue analysis cannot capture the molecular heterogeneity within distinct neuronal and glial populations, limiting insight into how NME and gut dysbiosis orchestrate long-lasting changes at the cellular level. A high-resolution approach is required to delineate the specific contributions of midbrain cell types to the enduring neuroinflammatory and sensory consequences of neonatal opioid exposure. To fill this knowledge gap, we employed single-cell RNA sequencing (scRNA-seq) to interrogate the transcriptional landscape of over 107,427 individual midbrain cells from adolescent mice exposed to NME, with or without *B. infantis* treatment, alongside saline controls. This high-resolution approach enables the identification of specific cell types affected by NME and reveals disrupted signaling pathways within each population. Moreover, it provides insight into how microbiome-based interventions may restore gene expression patterns associated with pain hypersensitivity and neurodevelopmental dysfunction following early opioid exposure.

## Materials and methods

### Animals

Adult male and female C57BL/6J mice (8–10 weeks old; The Jackson Laboratory, Bar Harbor, ME, USA) were used to generate experimental litters. Breeding pairs were housed in a temperature- and humidity-controlled environment under a 12-h light/dark cycle with ad libitum access to food and water. No more than five animals were housed per cage, and bedding was replaced regularly. Experimental groups were housed and handled separately to prevent cross-contamination of microbiota. Sample size was determined by power analysis (80% power, α = 0.05) based on preliminary data, indicating a minimum of five animals per group. Animals were euthanized using CO inhalation prior to tissue collection. All procedures were approved by the University of Miami Institutional Animal Care and Use Committee (IACUC) and conformed to NIH guidelines.

### Neonatal morphine exposure

Nulliparous female mice (10–12 weeks old) were housed with age-matched males (4 females:1 male) for timed mating. Pregnancy was confirmed by daily monitoring and pregnant dams were singly housed. The day of birth was designated as postnatal day 0 (P0). On postnatal days 6 or 7, pups were randomly assigned to receive either morphine sulfate (5 mg/kg/day, subcutaneous; NME group) or sterile saline (control group) for five consecutive days. All injections were administered in a total volume of 10 μL once daily. The morphine dose was selected based on prior reports demonstrating analgesic efficacy at 3 mg/kg, while avoiding excessive exposure associated with higher doses (10–15 mg/kg). During treatment, pups were briefly separated from the dam, weighed, and maintained on a heated pad to preserve body temperature. Pups were returned to the dam immediately following each procedure. After treatment, litters remained undisturbed with the dam until weaning at P21.

### Neonatal probiotic administration

Probiotic or vehicle treatments were administered to neonatal mice concurrently with neonatal morphine exposure (NME). *Bifidobacterium longum* subspecies *infantis* (*B. infantis*) was obtained in lyophilized powder form and reconstituted in sterile water. The solution was prepared to deliver a dose of up to 1 × 10 colony-forming units (CFU) per day, consistent with previously published protocols^7, 8^. Neonatal mice received either the *B. infantis* solution or vehicle (sterile water) via oral administration once daily for five consecutive days, delivered as droplets directly into the mouth. Probiotic treatment was administered during the same postnatal period as morphine or saline exposure. Upon weaning, pups were sexed and group-housed until adolescence (4–5 weeks of age). Female adolescents were used to collect the midbrain tissues.

### Tissue preparation

Tissue samples were processed following the 10x Genomics Single Cell Gene Expression Flex protocol. Briefly, dissected tissues were weighed to determine the required volume of Fixation Buffer B (1 mL per 25 mg of tissue). Tissues were finely minced on a pre-chilled glass petri dish maintained on ice to facilitate pipetting through a 1.5 mm wide-bore pipette tip. The minced tissue was resuspended in Fixation Buffer B and triturated gently using a wide-bore pipette. The suspension was transferred to a 2 mL centrifuge tube and triturated further until uniformly fragmented. Samples were incubated at 4°C for 16–24 hours to allow adequate fixation.

After fixation, samples were centrifuged at 850 × g for 5 minutes at 4°C. The supernatant was discarded, and the tissue pellet was washed with 2 mL of chilled phosphate-buffered saline (PBS). Following centrifugation at 850 × g for 5 minutes at room temperature, the supernatant was removed, and the tissue pellet was resuspended in 1 mL of Quenching Buffer B on ice. The sample was centrifuged again at 850 × g for 5 minutes, and the supernatant was discarded. The fixed tissue pellet was then prepared for downstream dissociation.

### Single-cell suspension generation

Fixed tissue pellets were dissociated by adding 2 mL of pre-warmed (37°C) Dissociation Solution. Samples were processed using an Octo Dissociator to ensure consistent and efficient dissociation. The resulting cell suspension was filtered through a 30 µm cell strainer to remove debris and undissociated tissue fragments. To maximize cell recovery, the filter was rinsed with an additional 2 mL of PBS. The filtrate was centrifuged at 850 × g for 5 minutes, and the resulting cell pellet was resuspended in 1 mL of chilled Quenching Buffer B. Cell concentration was determined using a hemocytometer or an automated cell counter (Countess II/3 FL). Fixed single-cell suspensions were processed immediately for downstream partitioning and library preparation using the 10x Genomics Chromium GEM-X Flex Single Cell Gene Expression workflow or stored in appropriate buffers according to manufacturer recommendations.

### 10x genomics single-cell RNA sequencing

10x Genomics Chromium Fixed RNA Profiling was performed by the Hussman Institute for Human Genomics (HIHG) at the University of Miami. Cell suspensions were assessed using the Nexcelom Cellometer K2. The suspensions were prepped following the Chromium Fixed RNA Profiling Reagent Kits for Multiplexed Samples User Guide.

Briefly, the initial step involved attaching Probe Barcodes to fixed samples through hybridization. Subsequently, these tagged samples were divided into extremely small, nanoliter-sized compartments called Gel Beads-in-emulsion (GEMs) using a microfluidic chip. To uniquely identify the molecules within each GEM, a selection from a large set of approximately 737,000 distinct 10x GEM Barcodes was introduced separately. Inside these GEMs, probes were joined together (ligated) with a 10x GEM Barcode, ensuring that all ligated probes within the same GEM carried the same barcode. These barcoded and ligated probes then underwent a bulk pre-amplification process, followed by the creation of gene expression libraries for sequencing by Illumina NovaSeq X Plus.

### scRNA-seq raw data processing

Demultiplexing, barcode processing, and single-cell gene expression matrix generation were performed using the Cell Ranger multi pipeline (v.8.0.1, 10X Genomics) with the Chromium Mouse Transcriptome Probe Set v1.0.1 and default parameters. Our dataset, comprising 13 samples, initially contained 121,856 cells. Initial processing and quality control of the scRNA-seq data were conducted using the Seurat package (v.5.0)^9^ in R (v.4.1.0) and RStudio (v.2022.05) implemented in Pegasus Cluster (CentOS 7) hosted by Institute for Data Science & Computing at University of Miami. Doublet cells were individually removed for each sample using scDblFinder package^10^. Additionally, we filtered the cells with the following parameters: maximum percentage of mitochondrial RNA = 10, minimum number of nFeature_RNA = 250 and minimum number of nCount_RNA = 500, maximum number of nFeature_RNA = 8,000 and maximum number of nCount_RNA = 35,000. Following quality control, 107,427 cells from 13 samples passed the initial QC and were retained for downstream analysis. These cells were then merged and integrated using the reciprocal PCA (RPCA) method implemented in the IntegrateLayers function of the Seurat package^9^, to minimize potential batch effect across experimental conditions. The integrated data were subsequently clustered using the FindClusters function with a resolution of 0.8, resulting in 48 initial clusters. Cell type annotation was performed in R using the sc-type pipeline^11^ according to the steps outlined https://github.com/IanevskiAleksandr/sc-type. Their built-in cell marker database was used, which details can be accessed at https://sctype.app/database.php. After the cell type annotation, 13 cell types were retained for downstream analysis.

### Identification of differential expressed genes (DEGs) across conditions, pathway analysis, upstream regulator analysis and cell–cell communication analysis

Differentially expressed genes (DEGs) across conditions for all cell types were identified using the FindMarkers function in Seurat, employing a built-in Wilcoxon test with Benjamini-Hochberg correction of p-values. Genes with an absolute Log2 fold change (|Log2FC|) > 0.25 and an adjusted p-value < 0.005 were considered statistically significant. These DEGs were then analyzed using QIAGEN Ingenuity Pathway Analysis (IPA)^12^ for canonical pathways enrichment analysis. Fisher’s exact test was used in these analyses to identify signaling and metabolic pathways significantly associated with the DEGs (p < 0.05). Upstream regulator analysis was also performed using the DEGs within the IPA software, with upstream regulators having a p-value < 0.05 considered significant. Cell–cell communication analysis was performed by CellChat R packages (v 2.1.2)^13, 14^. Replicate files from each condition were aggregated prior to CellChat object creation. The default CellChatDB for mouse were used, including secrete signaling interactions, extracellular matrix (ECM)-receptor interactions, cell-cell contact interactions and non-protein signaling (metabolic and synaptic signaling).

## Results

### Neonatal morphine exposure and probiotic supplementation did not alter the overall cellular composition of the adolescent mouse midbrain

This study investigated the long-term effects of neonatal morphine exposure (NME) on cell-type-specific transcriptional changes in the adolescent mouse midbrain using a murine model. Newborn mice were administered morphine at a concentration of 5 mg/kg/day or saline for five days (Figure 1A). A separate group of neonatally morphine-exposed mice received supplementation with the probiotic *B. infantis* (Figure 1A). Single-cell RNA sequencing (scRNA-seq) libraries were generated from 13 mouse midbrains. Midbrains were dissected from mice in the saline (Sal), morphine (Mor), and morphine plus probiotic (Mor+Pro) groups. These tissues were then dissociated into single-cell suspensions and fixed with formaldehyde.

**Figure 1.**
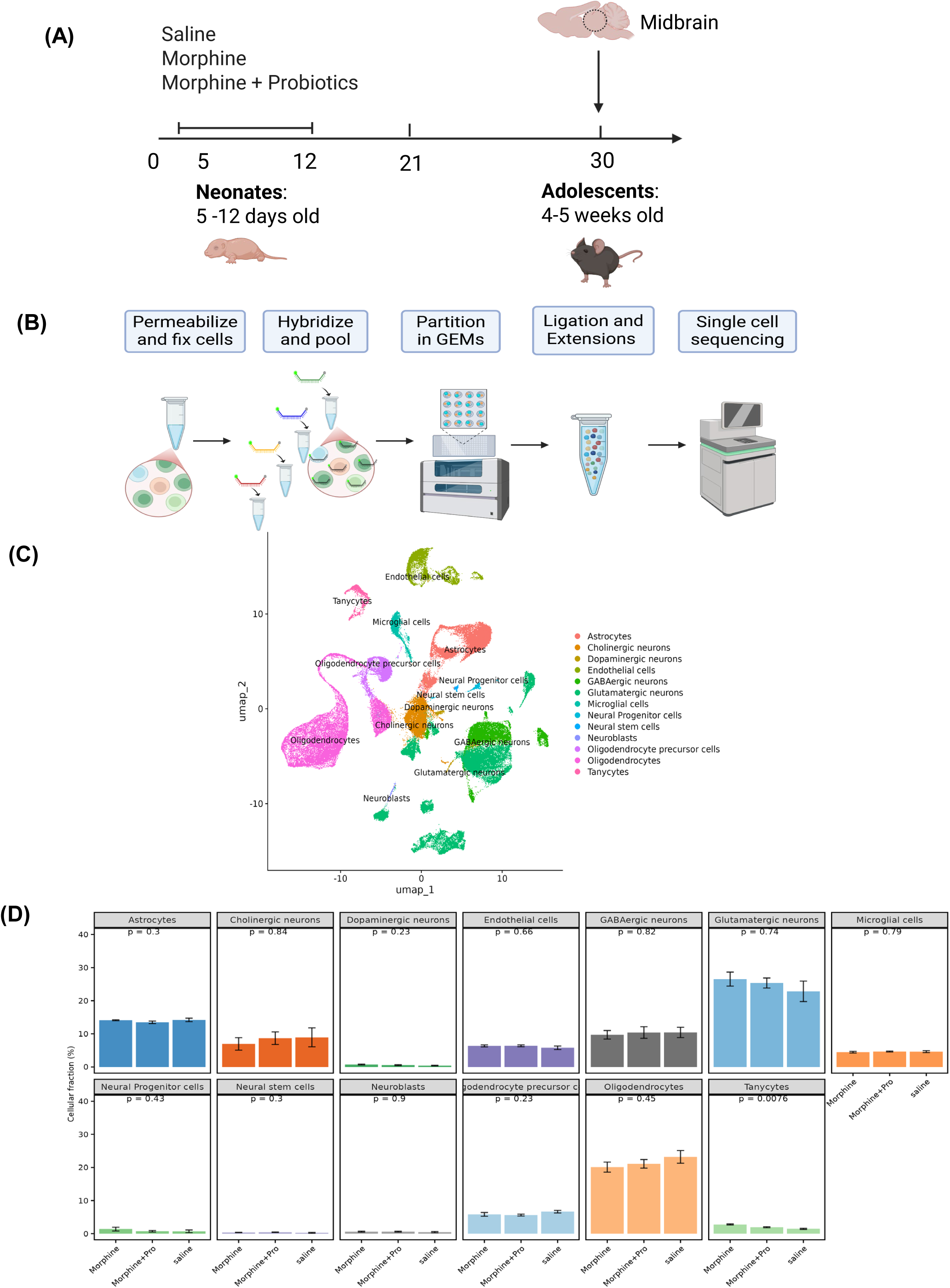
Consistent proportions of cell types across treatment groups. (A-B) Animal treatment design and single cell workflow for 13 samples. (C). Uniform Manifold Approximation and Projection (UMAP) plot showing the identified 13 major cell types, for total of n = 107,427 cells. D. Box plot of the 13 major cell types in each sample, split by treatment groups. See also Figure S1.

Multiplexing antibody barcodes facilitated the sequencing of multiple samples per channel. Cells were captured using the 10X Chromium platform, followed by Illumina sequencing, read alignment, and processing with 10X Cellranger (Figure 1B). Following doublet removal, quality control filtering, and data integration (see Methods and Figure S1B-C), a total of 107,427 single cells with complete transcriptional profiles were obtained. These cells were subsequently clustered using Seurat and assigned to 13 distinct cell types using the Sc-type pipeline, which integrates data from the CellMarker^15^ database and PanglaoDB^16^ (Figure 1C).

Next, we investigated whether neonatal morphine exposure altered the proportions of different cell types in the adolescent midbrain. Across all three experimental groups, neurons constituted approximately 40% of the total cell population. Oligodendrocytes and oligodendrocyte precursor cells (OPCs) combined accounted for 26%, followed by astrocytes (14%), epithelial cells (6%), microglia (5%), with the remaining cell types each representing less than 5% of the total cell count (Figure S1A). A Kruskal-Wallis test was performed to determine if any cell type proportions differed significantly among the three conditions. This analysis revealed no significant differences (p > 0.05) in the proportions of most cell types across the groups, with the exception of tanycytes (p = 0.008) (Figure 1D). Consistent with this finding, Uniform Manifold Approximation and Projection (UMAP) plots showed highly similar distributions of cell types across the Sal, Mor, and Mor+Pro groups (Figure S1D).

### Neonatal morphine exposure altered the expression of thousands of gene transcripts, in particular HOX genes, in several cells of the adolescent midbrain

We first analyzed cell-type-specific differentially expressed genes (DEGs) to investigate transcriptional changes induced by morphine (Mor vs. Sal). Under the morphine condition, each cell type displayed distinct changes, as illustrated in the combined volcano plots (Figure 2A). Specifically, more than one thousand DEGs were identified in glutamatergic and GABAergic neurons, oligodendrocytes, and astrocytes. In contrast, fewer than five hundred DEGs were observed in microglia, OPCs, and cholinergic neurons (Figure 2B). An upset plot (Figure 2B) revealed that most DEGs were unique to each cell type. Glutamatergic and GABAergic neurons, oligodendrocytes, and astrocytes each had over 500 unique DEGs, whereas microglia, OPCs, and cholinergic neurons had fewer than 200. Notably, a set of HOX genes exhibited significant downregulation with the lowest Log2 fold change in glutamatergic and GABAergic neurons, oligodendrocytes, and astrocytes under morphine conditions (Figure 2A).

**Figure 2.**
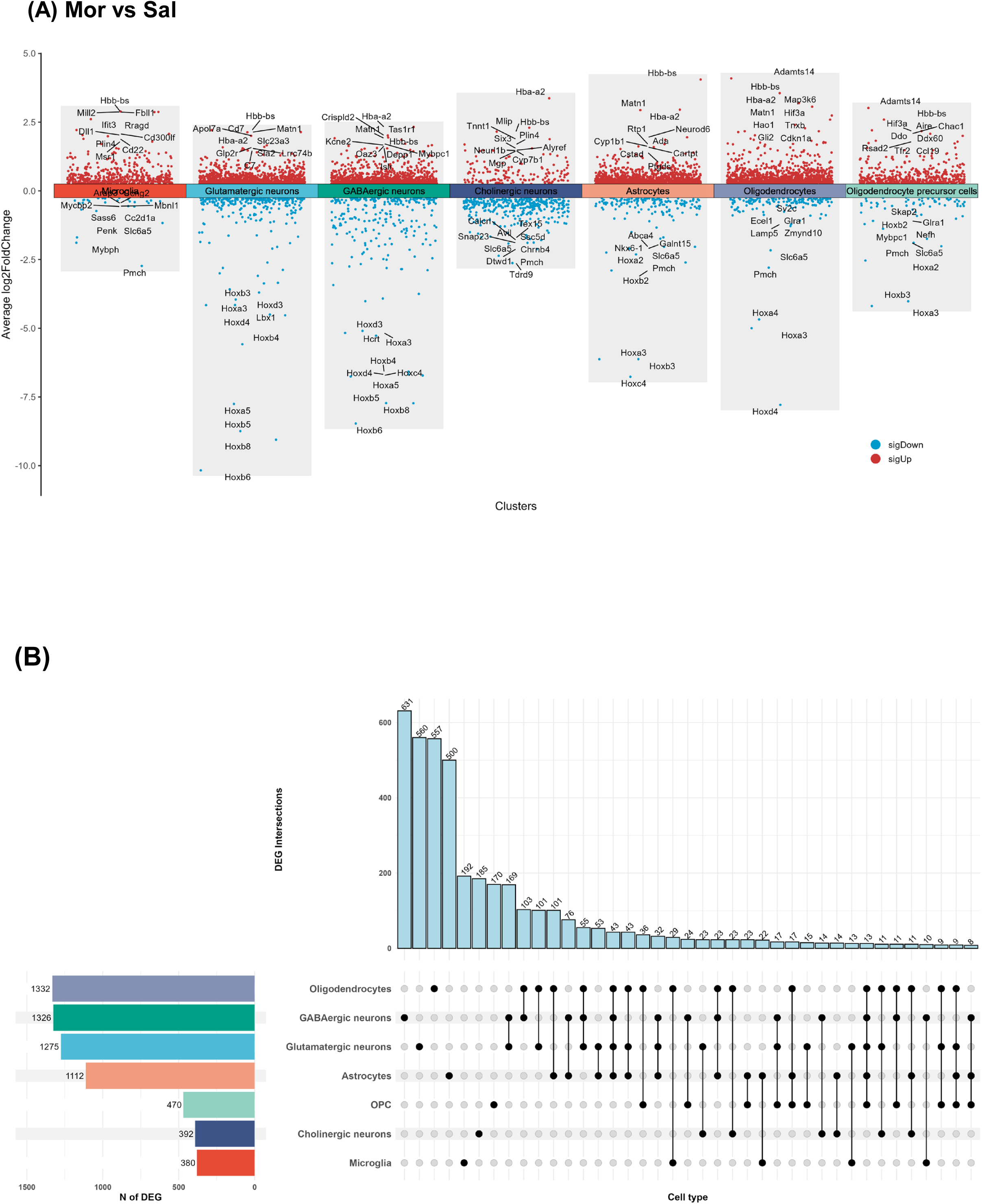
Differentially expressed genes (DEGs) under Mor vs Sal. (A) Combined volcano plots Representation of differentially expressed genes (DEGs) across cell types. (B) UpSet diagram represents counts of shared DEGs across cell types and number (*N*) of DEGs for each cell type. Genes with |Log2FC| > 0.25 and adjusted p value < 0.005 are counted as significantly changed. See also Table S1.

### Neonatal morphine exposure induced canonical pathways changes including neurotransmitter signaling and cellular growth and proliferation in the midbrain

To further investigate genome-wide alterations in gene transcripts, we performed a QIAGEN IPA canonical pathway enrichment analysis, focusing on cell types with the highest number of DEGs. We then generated enrichment bubble plots for glutamatergic neurons, GABAergic neurons, astrocytes, and oligodendrocytes (Figure 3). Under morphine conditions (Mor vs. Sal), several canonical pathways, including gene transcription, signal transduction, cell growth and proliferation, and neurotransmitter signaling, were significantly upregulated in glutamatergic neurons (Figure 3A). Similarly, in GABAergic neurons, signal transduction, cellular growth and proliferation, and neurotransmitter signaling were upregulated under morphine conditions (Figure 3B). In astrocytes, under morphine conditions, we observed increased metabolism, neurotransmitter signaling, organismal growth and development, and disease-specific pathways (Figure 3C). Under morphine conditions, oligodendrocytes showed increased neurotransmitter signaling, biosynthesis, disease pathways, and cellular stress and injury (Figure 3D).

**Figure 3.**
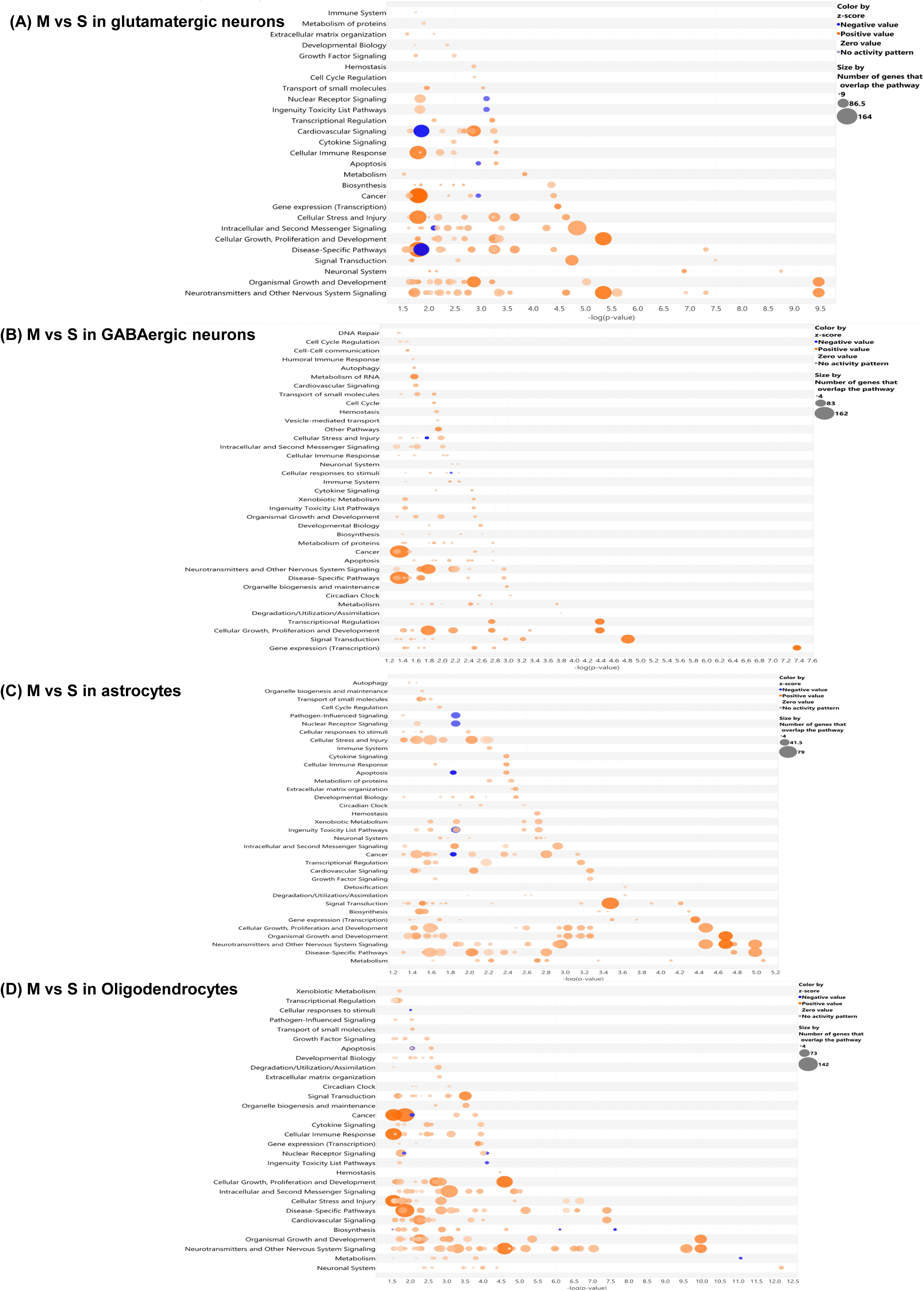
IPA analysis with canonical pathways under Mor vs Sal of (A) canonical pathways under Mor vs Sal of Glutamatergic neurons. (B) canonical pathways under Mor vs Sal of GABAergic neurons. (C) canonical pathways under Mor vs Sal of Astrocytes. (D) canonical pathways under Mor vs Sal of Oligodendrocytes. M vs S, morphine vs saline. See also Table S2.

### Probiotic supplementation reversed the expression of HOX genes in specific cell types induced by neonatal morphine exposure in the midbrain

Previous studies from our group have shown that neonatal probiotic intervention with *B. infantis* reversed the pain hypersensitivity and mitigated NME-induced transcriptional changes in the midbrain by attenuating gut dysbiosis^4^. Therefore, to assess the impact of gut microbiome-targeted intervention, we then analyzed cell-type-specific DEGs under probiotic supplementation (Mor+Pro vs. Mor) and compared them to the saline control (Mor+Pro vs. Sal). Interestingly, HOX genes downregulated under morphine conditions showed significant upregulation with probiotic supplementation. However, this reversal was specific to glutamatergic and GABAergic neurons, oligodendrocytes, astrocytes, and OPCs (Figure 4, S2). In glutamatergic and GABAergic neurons, probiotics supplementation led to a 64-fold increase (6 units of Log2 fold change increase) of HOX genes (Figure 4A, B). However, when compared to saline control, HOX genes were still downregulated with a log2 fold change of -4, suggesting the probiotics supplementation did not completely reverse the downregulation of HOX genes. In astrocytes and oligodendrocytes, probiotics supplementation similarly caused a 64-fold increase in gene expression (Figure 4C, D). Interestingly, when compared to saline control, HOX genes were no longer among the most regulated genes, suggesting that probiotics supplementation successfully reversed the downregulation of HOX genes. In OPCs, probiotics supplementation induced a smaller increase of HOX genes and was able to reverse the downregulation of HOX genes from morphine (Figure S2A). In contrast, HOX genes were not downregulated in microglia and cholinergic neurons under morphine conditions nor did probiotics supplementation lead to increased HOX gene expression (Figure S2B, C).

**Figure 4.**
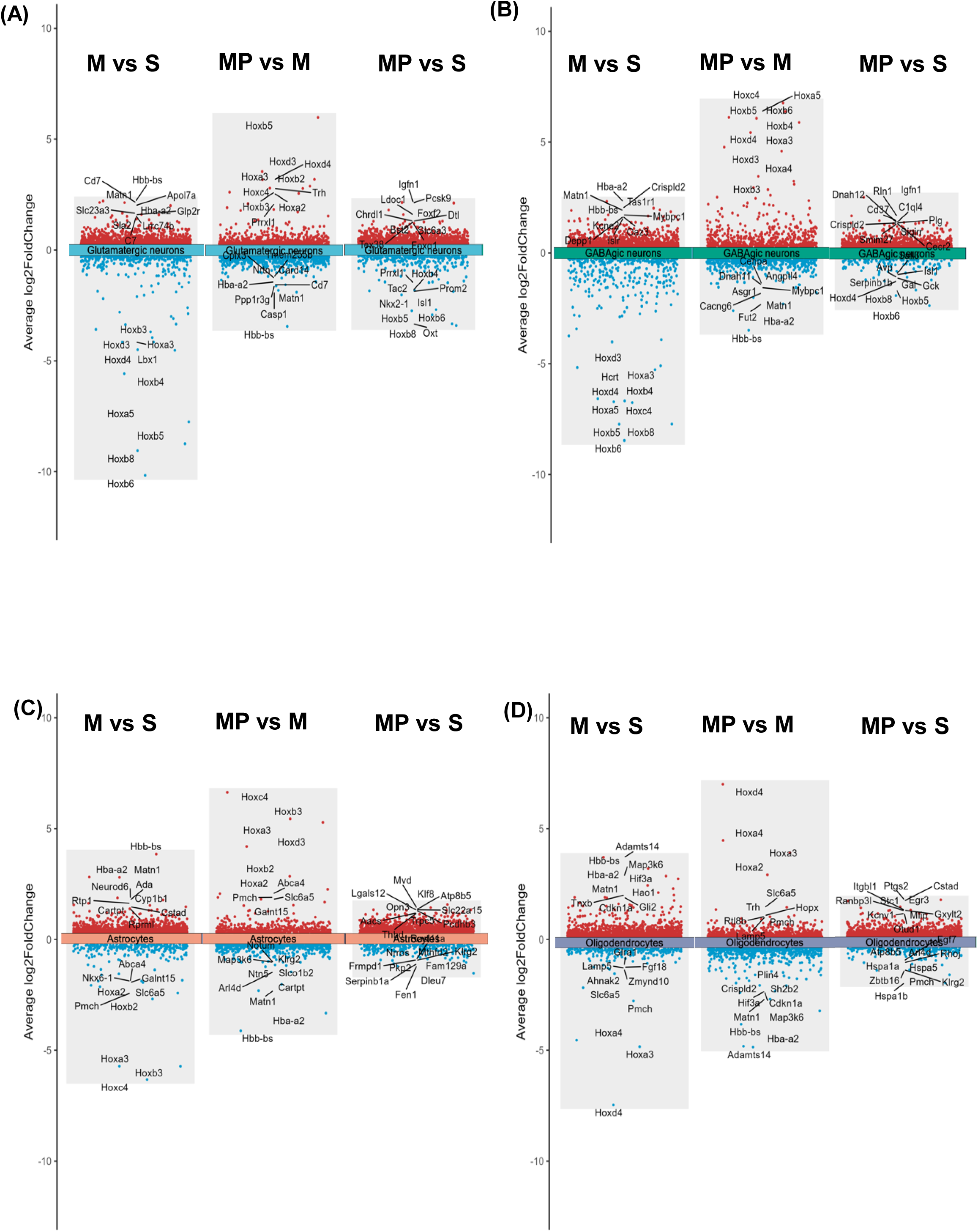
Side-by-side DEGs comparisons of Glutamatergic neuron (A), GABAergic neuron (B), astrocyte (C), and oligodendrocytes (D) in Mor vs Sal, Mor+Pro vs Mor and Mor+Pro vs Sal conditions. Genes with |Log2FC| > 0.25 and adjusted p-value < 0.05 are counted as significantly changed. M vs S, morphine vs saline. MP+M, morphine +probiotics vs morphine. MP vs S, morphine+probiotics vs saline. See also Figure S2 and Table S1.

### Probiotic supplementation reversed canonical pathways changes that were induced by neonatal morphine exposure in the midbrain

To further investigate the impact of probiotics supplementation on canonical pathways, we performed a QIAGEN IPA canonical pathway enrichment analysis under probiotic supplementation (Mor+Pro vs. Mor), focusing on glutamatergic neurons, GABAergic neurons, astrocytes, and oligodendrocytes (Figure 5). Under probiotic supplementation (Mor+Pro vs. Mor), the majority of pathways that were upregulated by neonatal morphine exposure were downregulated in glutamatergic neurons (Figure 5A), with neurotransmitter signaling, cellular growth and development, and cellular immune response among the most significantly downregulated. Similarly, in GABAergic neurons, probiotic supplementation led to the downregulation of neurotransmitter signaling, organismal growth and development, transcription regulation, and cell stress and injury (Figure 5B). In astrocytes, probiotic supplementation resulted in the downregulation of disease-specific pathways, neurotransmitter signaling, and cell stress and injury in astrocytes (Figure 5C). Probiotic supplementation in oligodendrocytes led to decreased biosynthesis, intracellular signaling, degradation utilization, and gene transcription (Figure 5D).

**Figure 5.**
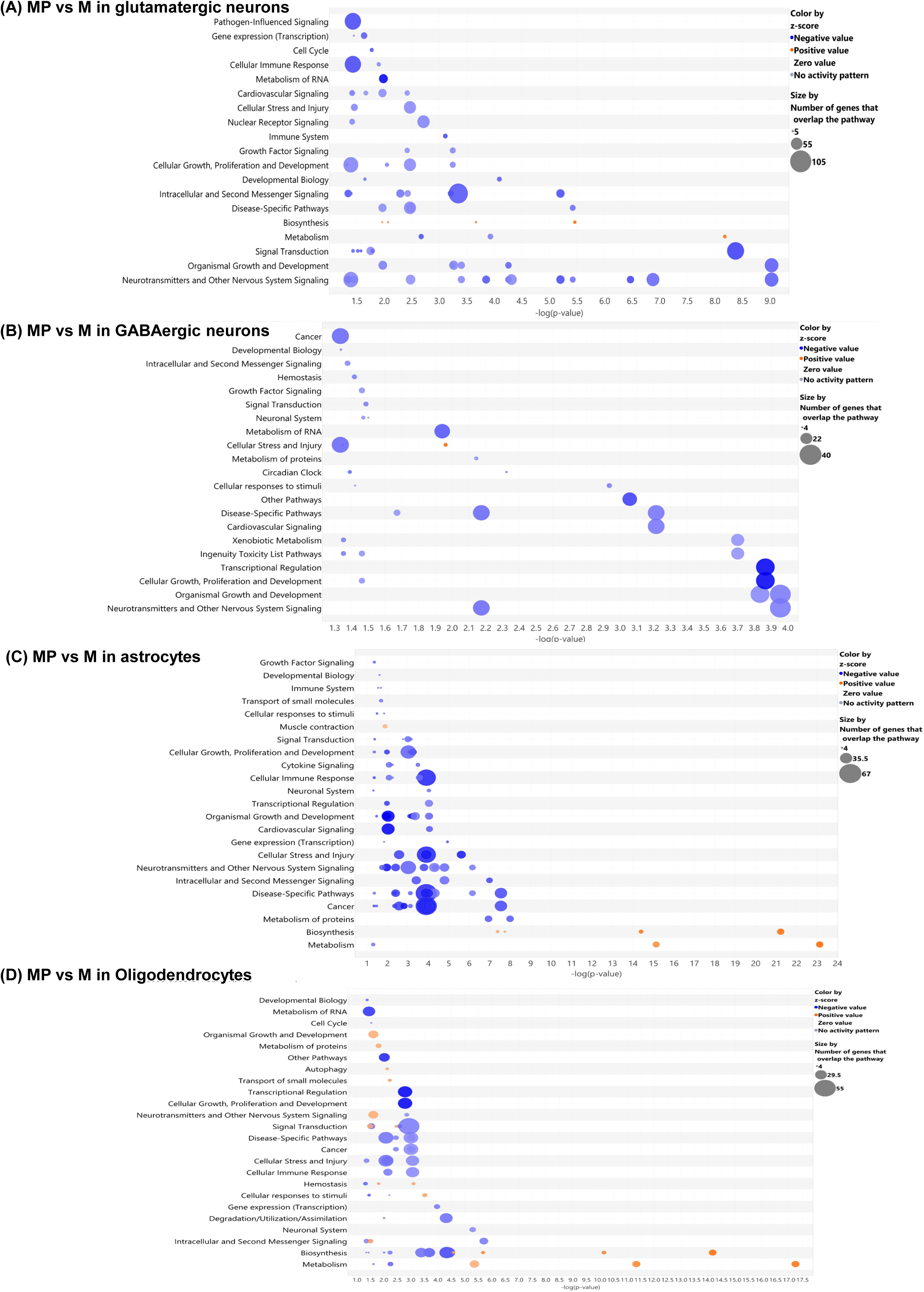
IPA analysis with canonical pathways under Mor+Pro vs Mor of (A) canonical pathways under Mor+Pro vs Mor of Glutamatergic neurons. (B) canonical pathways under Mor+Pro vs Mor of GABAergic neurons. (C) canonical pathways under Mor+Pro vs Mor of Astrocytes. (D) canonical pathways under Mor+Pro vs Mor of Oligodendrocytes. MP vs M, morphine +probiotics vs morphine. See also Table S2.

### Neonatal morphine exposure enhanced neuropathic pain pathways, while probiotic supplementation attenuated them

Previous studies from our group have shown that neonatal morphine exposure induces prolonged pain hypersensitivity during adolescence^4^, Therefore, we investigated how pain-related canonical pathways were affected by morphine and probiotic supplementation in specific cell types. Glutamatergic neurons play a significant role in neuropathic pain. Under morphine conditions (Mor vs. Sal), these neurons exhibited enhanced overall neuropathic pain pathways (Figure 6A). Specifically, multiple subunits of Metabotropic Glutamate Receptors (mGluR-GRM), Ionotropic Glutamate Receptors AMPA type (GRIA – AMPA), and Ionotropic Glutamate Receptors NMDA type (NMDA) were upregulated (Figure S4A). However, probiotic supplementation (Mor+Pro vs. Mor) attenuated these pain pathways in glutamatergic neurons (Figure 6B). Most glutamate transporter genes were downregulated including mGluR-GRM and NMDA receptors (Figure S4B). No significant changes in pain pathways were observed in inhibitory GABAergic neurons (data not shown). Astrocytes also modulate neuropathic pain through interactions with neurons and other glial cells. Similar to glutamatergic neurons, astrocytes showed enhanced neuropathic pain pathways under morphine conditions (Figure 6C) and decreased pain pathways with probiotic supplementation (Figure 6D). Similarly, mGluR-GRM and NMDA receptors were upregulated with morphine (Figure S4C) and downregulated with probiotics supplementation (Figure S4D). Oligodendrocytes displayed a similar trend of increased and decreased neuropathic pain pathways with morphine exposure and probiotic supplementation, respectively (Figure S3, S4E-F).

**Figure 6.**
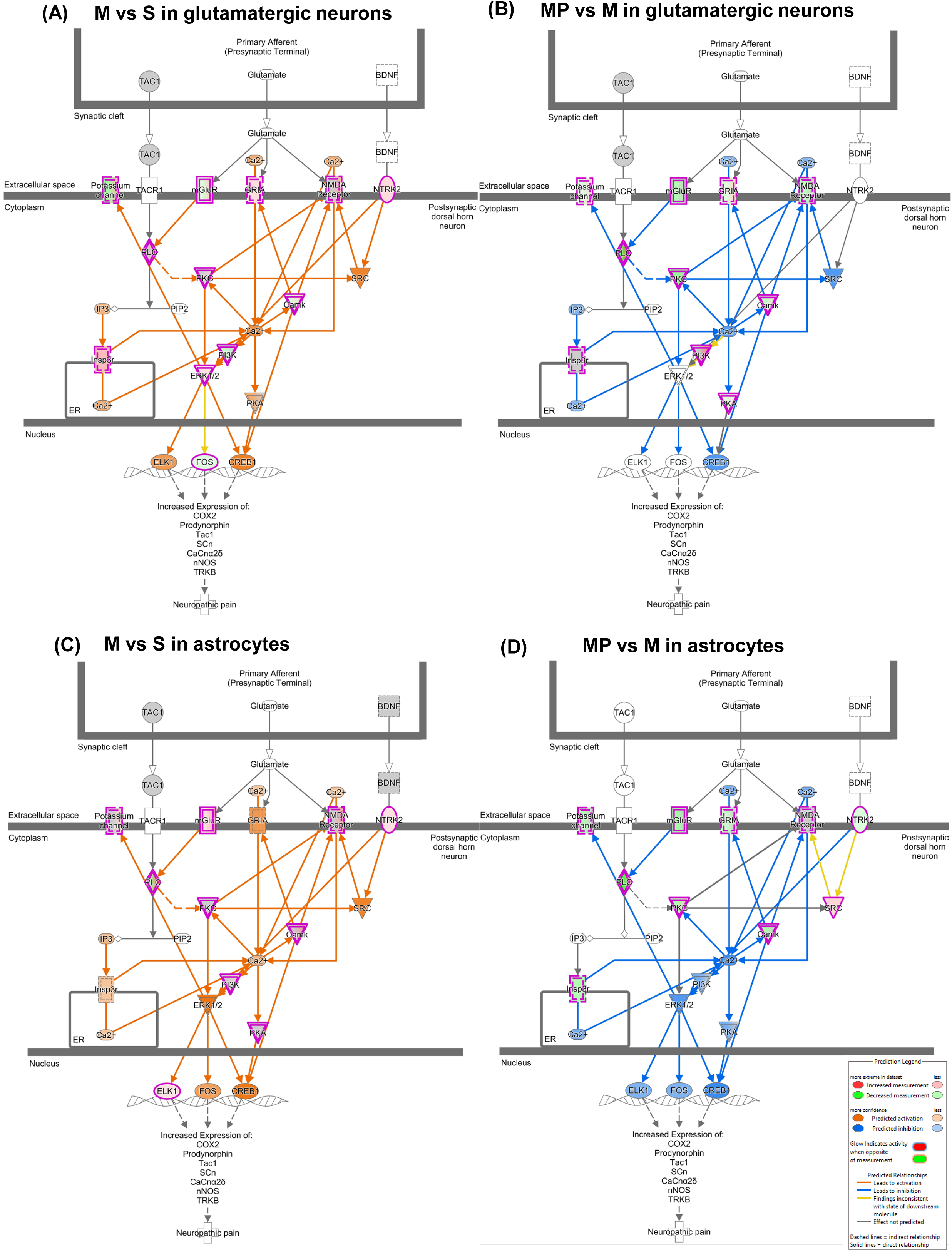
Neuropathic pain pathways of Glutamatergic neurons under Mor vs Sal (A) and Mor+Pro vs Mor (B) conditions. Astrocytes under Mor vs Sal (C) and Mor+Pro vs Mor (D) conditions. M vs S, morphine vs saline. MP+M, morphine +probiotics vs morphine. See also Figure S3, S4.

### Neonatal morphine exposure increased cell-cell communication between glutamatergic neurons, astrocytes, and oligodendrocytes, and probiotic supplementation attenuated this communication

Next, we analyzed cell-cell communication using CellChat (see Methods) among glutamatergic neurons, astrocytes, and oligodendrocytes, to investigate how cell communication was affected by morphine and probiotic supplementation. We first compared the number and strength of these interactions between glutamatergic neurons, astrocytes, and oligodendrocytes to identify significant changes under morphine and probiotic conditions. Based on circle plots (Figure 7A, S5A), both the number and strength of interactions from glutamatergic neurons to glial cells increased under morphine conditions (Mor vs. Sal). Similarly, the number and strength of interactions from glial cells to glutamatergic neurons also increased (Figure 7A, S5A), indicating bidirectional enhancement of cell-cell communication with morphine exposure. Under probiotic supplementation (Mor+Pro vs. Mor), we observed a reversal of these trends; both the number and strength of interactions from glutamatergic neurons to glial cells decreased (Figure 7B, S5B), as did the communication from glial cells to glutamatergic neurons (Figure 7B, S5B), suggesting that probiotic treatment reduced bidirectional communication between these cell types. Next, we conducted information flow analysis to identify signaling pathways altered by the different conditions, by calculating the sum of communication probabilities among all ligand-receptor pairs within the inferred network. Under morphine conditions (Mor vs. Sal), glutamate and neurexin (NRXN) pathways showed increased overall information flow from glutamatergic neurons to astrocytes and oligodendrocytes (Figure 7C). However, under probiotic supplementation (Mor+Pro vs. Mor), these pathways exhibited reduced information flow from glutamatergic neurons to astrocytes and oligodendrocytes (Figure 7D). Similarly, morphine exposure increased information flow of glutamate and NRXN pathways from astrocytes and oligodendrocytes to glutamatergic neurons (Figure S5C), while probiotic treatment decreased this information flow (Figure S5D).

**Figure 7.**
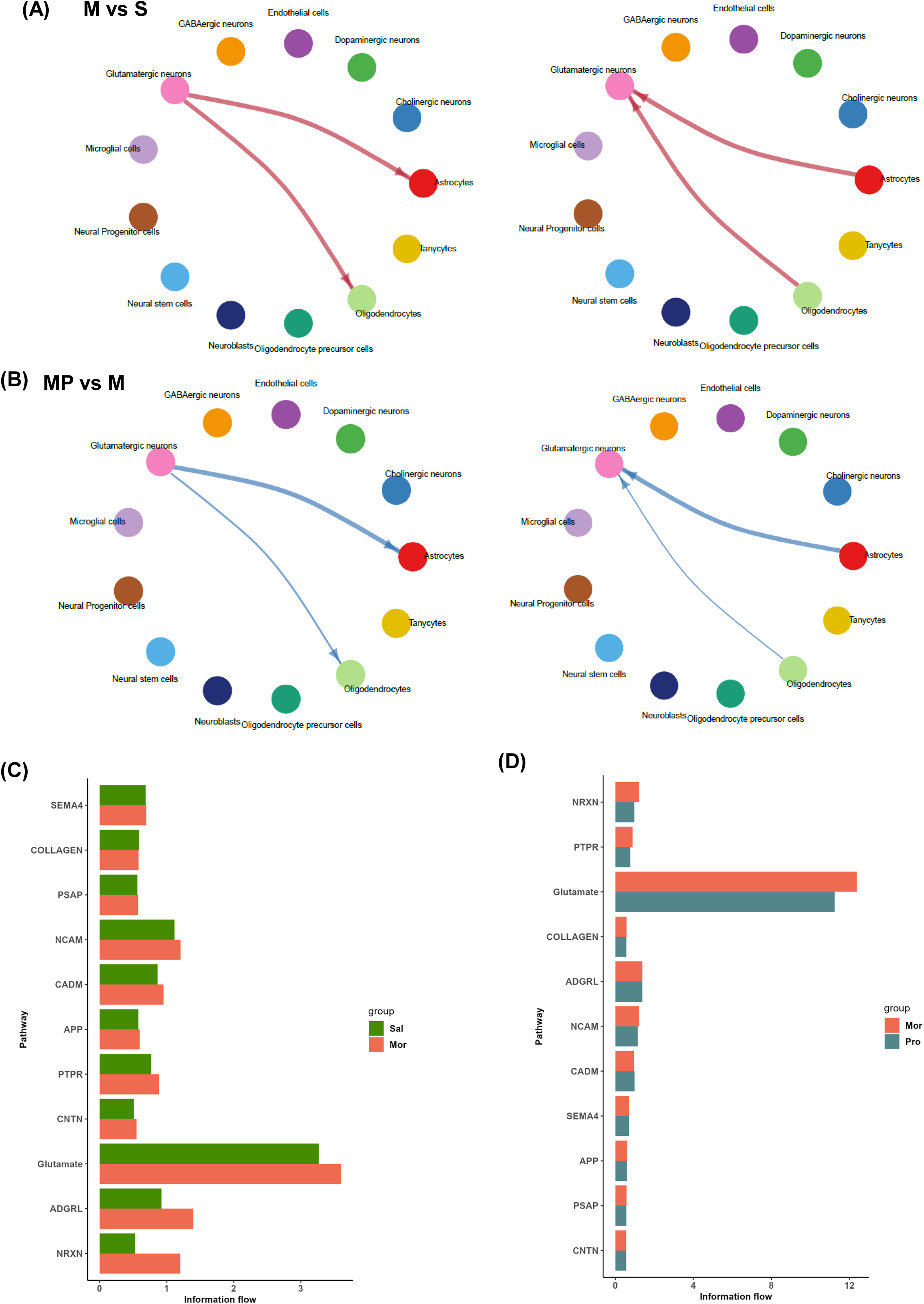
cell–cell communication among Glutamatergic neurons, astrocytes and oligodendrocytes. (A) Circle plot showing differential number of interactions between Glutamatergic neurons, astrocytes and oligodendrocytes under M vs S. Red (or blue) colored edges represent increased (or decreased) signaling in the comparison. (B) Circle plot showing differential number of interactions between Glutamatergic neurons, astrocytes and oligodendrocytes under MP+M. (C) Overall information flow of top signaling pathway from Glutamatergic neuron to astrocytes and oligodendrocytes under M vs S (D) Overall information flow of top signaling pathway from Glutamatergic neuron to astrocytes and oligodendrocytes under MP+M. M vs S, morphine vs saline. MP+M, morphine +probiotics vs morphine. See also Figure S5.

### Upstream regulator analysis indicated that retinoic acid receptors (RARs) were downregulated by neonatal morphine exposure and upregulated by probiotic supplementation

Finally, to investigate the potential regulatory mechanism, we performed IPA Upstream Regulator Analysis to identify potential upstream regulators based on the observed gene expression changes in our scRNA-seq dataset. Given the significant fluctuations in HOX gene expression observed in specific cell types under morphine and probiotic conditions (Figure 4), we focused our analysis on these genes to determine their regulatory relationships. Specifically, under morphine conditions (Mor vs. Sal), Retinoic acid receptors (RARs) alpha and gamma were predicted to be significantly downregulated in glutamatergic neurons (Figure 8A, S6A). Furthermore, RAR/retinoid X receptor (RXR) heterodimers also showed predicted lower expression under morphine conditions (Figure S6A). Conversely, RAR-alpha, RAR-gamma, and the RAR/RXR complex were all predicted to be significantly upregulated in glutamatergic neurons with probiotic supplementation (Mor+Pro vs. Mor) (Figure 8B, S6B). Similarly, in GABAergic neurons under morphine conditions, RAR-alpha and RAR-gamma were predicted to be significantly downregulated (Figure 8C, S6C). However, probiotic supplementation led to the predicted upregulation of RAR-alpha, RAR-gamma, and the RAR/RXR complex in GABAergic neurons (Figure 8D, S6D). In astrocytes under morphine conditions, only RAR-alpha was predicted to be significantly downregulated (Figure 8E). Probiotic supplementation, however, was associated with the predicted upregulation of RAR-alpha, RAR-beta, and RAR-gamma in astrocytes (Figure 8F, S7). Finally, in oligodendrocytes, RAR-alpha was the sole regulator predicted to be downregulated under morphine conditions and upregulated with probiotic supplementation (Figure 8G-H).

**Figure 8.**
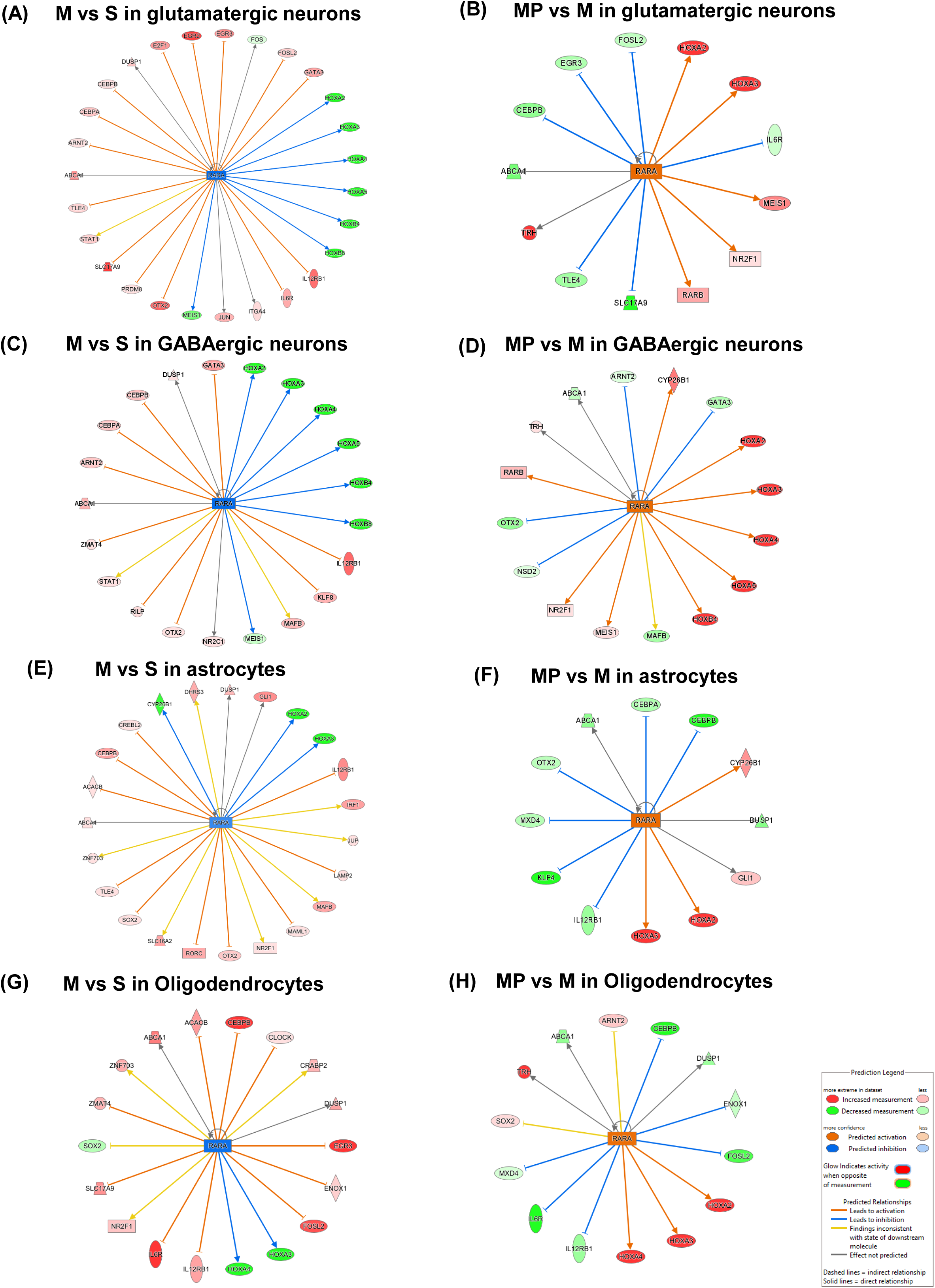
IPA Upstream Regulator Analysis of Glutamatergic neurons under Mor vs Sal (A) and Mor+Pro vs Mor (B) conditions, GABAergic neurons under Mor vs Sal (C) and Mor+Pro vs Mor (D) conditions, and Astrocytes under Mor vs Sal (E) and Mor+Pro vs Mor (F) conditions., Oligodendrocytes under Mor vs Sal (G) and Mor+Pro vs Mor (H) conditions. M vs S, morphine vs saline. MP+M, morphine +probiotics vs morphine. See also Figure S6, S7.

## Discussion

The study presents the first single-cell RNA sequencing dataset of the adolescent midbrain following neonatal morphine exposure and probiotic intervention. Using a murine model, we found that neonatal morphine exposure induced cell-type-specific changes in gene expression, disrupted canonical signaling pathways, and altered cell-cell communication in the adolescent midbrain. Notably, probiotic supplementation with *B. infantis* reversed many of these morphine-induced alterations in gene expression including those related to pain pathways, and cell-cell communication. These findings offer novel insights into the long-term impact of early opioid exposure on the brain and highlight potential mechanisms through which microbiome-targeted therapies may exert neuroprotective effects.

We found that neonatal morphine treatment did not significantly alter the proportions of most midbrain cell types. However, it disrupted gene expression profiles across multiple populations. Specifically, glutamatergic and GABAergic neurons, oligodendrocytes, and astrocytes had the greatest number of DEGs and unique DEGs. These findings are consistent with prior single-cell studies. For instance, a previous single-cell study showed that while proportion of each cell type was consistent, chronic morphine treatment did not influence the composition of the amygdala tissue, but did affect the transcriptional profiles of all cell types in mice amygdala tissue^17^. In that study, microglia, endothelial cells, neurons and oligodendrocyte progenitor cells had the greatest number of up- and down-regulated genes^17^. Similarly, single cell transcriptomics of postmortem midbrain tissue from individuals with a history of opioid use revealed that microglia, oligodendrocytes, OPCs, and astrocytes accounted for the majority of DEGs^18^. In that study as well, chronic opioid exposure and overdose were not associated with proportional shifts among the neuronal and glial constituents^18^. A separate study in human striatal tissue from individuals with opioid use disorder (OUD) found no significant differences in the proportions of neuronal and glial cell types, but identified 1,765 DEGs across cell types, with glial cells showing more transcriptional changes than neurons^19^. Taken together, these findings support a model in which opioid exposure alters cell-type-specific transcriptional programs without significantly affecting cellular composition. Variability across studies may reflect differences in opioid treatment regimens, single-cell sequencing techniques, or confounding factors in human subjects.

Additionally, we found that neonatal morphine exposure upregulated various canonical pathways related to neurotransmitter signaling, cellular growth, immune response, and disease-related pathways. These findings are consistent with our current understanding of opioid-induced neuroinflammation^20, 21^ and align with bulk RNA-sequencing data from our previous work using the same animal model^4^. Moreover, our results demonstrated that neurotransmitter signaling, cellular immune response, along other the pathways that were previously upregulated with neonatal morphine exposure were downregulated by probiotics intervention. Prior studies have reported similar pathway alterations in response to opioid treatment. For example, chronic morphine exposure in the amygdala was shown to upregulate pathways related to cell activation, immune and inflammatory responses, cytokine signaling, and metabolic processes^17^. In postmortem human ventral midbrain tissue, glial cell populations—including astrocytes, pericytes, microglia, and oligodendrocytes—exhibited strong upregulation of immune-related pathways, including interferon signaling, NF-κB activation, and cell motility^18^. In the striatum of individuals with opioid use disorder (OUD) and opioid-exposed rhesus macaques, neurons showed enrichment of pathways linked to neurodegeneration, interferon responses, and DNA damage, while glial cells displayed upregulation of neuroinflammation and synaptic signaling pathways^19^. Furthermore, a single-nucleus RNA-seq study of brain organoids derived from patients with OUD found that oxycodone induced type I interferon signaling in neurons and astrocytes, whereas buprenorphine activated the mTOR pathway in astrocytes^22^. However, these previous studies did not examine therapeutic interventions and thus provide no data on the potential modulatory effects of probiotics. Our findings fill this gap by demonstrating that microbiome-targeted treatment can reverse morphine-induced transcriptional and pathway-level changes.

One of the most striking findings from our study was the morphine-induced downregulation of homeobox (Hox) genes across multiple midbrain cell types. Hox genes are highly conserved developmental transcription factors that play essential roles in establishing positional identity, regional specification, tissue patterning, and cell differentiation^23, 24^. Although direct evidence linking Hox genes to opioid addiction is limited, several of these genes have been implicated in nociceptive circuit formation in the spinal cord, particularly relevant given our prior observation that neonatal morphine exposure (NME) leads to persistent pain hypersensitivity during adolescence^4^. Hox genes are critically involved in the formation of the hindbrain and spinal cord, where they regulate the identity and connectivity of spinal interneurons involved in pain processing. ^25, 26, 27, 28^. The expression of distinct combinations of Hox genes in the hindbrain rhombomeres is required to generate the precursors of visceral sensory interneurons^29^. Additionally, loss of *Hoxb8* in a mice model led to excessive lower back grooming, localized skin lesions in adult mutant mice, and attenuated response to nociceptive and thermal stimuli^30^. Importantly, in spinal cord circuits, several studies revealed connectivity defects upon Hox gene inactivation. For example, early or late genetic removal of *Hox5* in mice affects diaphragm innervation, demonstrating that *Hox5* genes are required for proper connectivity of phrenic motor neurons to premotor interneurons^31^. *Hoxc8* removal specifically in sensory neurons affects sensory-motor connectivity^32^ whereas motor neuron-specific depletion of *Hoxc8* affects forelimb muscle innervation^33^. These studies highlight that Hox genes are important in establishing and maintaining normal neuronal wiring and synaptic plasticity during the post-natal stages. Importantly, our data show that probiotic supplementation restored the expression of several Hox genes across diverse midbrain cell types, including glutamatergic and GABAergic neurons, oligodendrocytes, astrocytes, and OPCs. Given the role of Hox genes in specifying, differentiating, and refining synaptic connections in pain-related neurons, their reactivation may represent a key mechanism through which probiotics counteract morphine-induced neurodevelopmental disruption. These results not only reinforce the critical role of Hox gene expression in maintaining neuronal function but also suggest that microbiome-targeted interventions can reverse opioid-induced transcriptional dysregulation and support neurodevelopmental resilience.

In addition to the observed homeobox gene disruptions, IPA analysis of our single-cell transcriptomic data revealed a coordinated upregulation of the neuropathic pain signaling pathway across multiple midbrain cell types following neonatal morphine exposure. Specifically, morphine-treated mice exhibited increased expression of glutamate receptor genes, including metabotropic glutamate receptors (mGluRs), AMPA receptors (GRIA), and NMDA receptors (GRN), which are well-known mediators of synaptic excitability and plasticity^34^. These receptors are critical for activity-dependent potentiation in nociceptive circuits and have been strongly implicated in the pathogenesis of central sensitization and pain^35, 36, 37^. Additionally, morphine exposure led to increased expression of key neurotrophin-related molecules, such as the neurotrophin receptor *NTRK2* (TrkB) which mediates brain-derived neurotrophic factor (BDNF) signaling. This pathway is known to enhance neuronal excitability^38^. We also identified upregulation of multiple intracellular signaling cascades, including phospholipase C (PLC), protein kinase C (PKC), calmodulin-dependent kinase (CaMK), phosphoinositide 3-kinase (PI3K), and extracellular signal-regulated kinases 1/2 (ERK1/2), all of which converge on transcriptional regulators like *CREB1* and *ELK1* to promote the long-term transcriptional programs sustaining chronic pain states^39, 40, 41^. These changes were accompanied by predicted increases in intracellular calcium signaling (*Ca²* , *IP3*, *IP3R*) and kinase activity (*SRC*, *PKA*), further indicating a morphine-induced shift toward hyperexcitability and maladaptive plasticity. Remarkably, probiotic supplementation reversed the expression direction of nearly all of these genes and predicted signaling events in multiple cell types, including glutamatergic neurons, astrocytes, and oligodendrocyte lineage cells. This reversal suggests that the probiotic not only mitigates transcriptional dysregulation associated with pain sensitization but may also restore homeostatic signaling in critical neurodevelopmental and neuromodulatory pathways. These findings are consistent with prior studies demonstrating the role of opioid-induced microbial dysbiosis in morphine-associated comorbidities, and the potential of probiotics to ameliorate such effects^4, 42, 43^. For example, probiotic VSL#3, containing several *Bifidobacterium* and *Lactobacillus* species, has been shown to prevent the development of morphine tolerance by restoring microbial balance^42^. Similarly, Cai et al. reported that gut microbiota from fibromyalgia patients induced pain behavior in mice, which was reversed by microbiota from healthy individuals, suggesting a causal role for microbiota in widespread pain^44^. In our previous study, we demonstrated that *B. infantis* supplementation in neonates mitigated pain hypersensitivity by restoring gut microbial composition and reducing systemic inflammation.^4^ The present study extends those findings by showing that *B. infantis* can also rescue morphine-induced transcriptional disruptions at the single-cell level. Together, these findings position probiotic intervention as a promising strategy to attenuate early-life morphine-induced molecular reprogramming that contributes to neurodevelopment deficits. Moreover, our cell-cell communication analysis demonstrated that there was a bidirectional increase in communications among glutamatergic neurons, astrocytes, and oligodendrocytes under morphine conditions. Particularly, glutamate pathway communication which was involved in pain signaling was among the most increased pathway under the morphine condition. Probiotics supplementations were able to decrease the bidirectional communication among cells as well as glutamate pathway. The cell-cell communication analysis results, along with pain pathway analysis, corroborate with our previous study that neonatal supplementation with the *probiotic B. infantis* prevented the onset of pain hypersensitivity^4^.

Finally, our upstream regulator analysis identified RARs and RXRs can be a potential regulator during neonatal morphine exposure and probiotics supplementation. Retinoic acid receptors (RARs) are a type of nuclear receptor that act as ligand-activated transcription factors. They are activated by all-trans-retinoic acid (ATRA), a derivative of vitamin A. There are three main types of RARs of RXRs: RAR-alpha, RAR-beta, and RAR-gamma. RXRs often form heterodimers with other nuclear receptors, including RARs, to regulate gene expression^45^. A multi-omics study on morphine effects on mice reported that all-trans-retinoic acid (atRA) were depleted in the in ileal luminal metabolome of morphine group^46^. Further, their bulk-RNA sequencing on ileum tissue result showed a decrease in retinol metabolism as well as decrease in Retinol Dehydrogenase 7 (Rdh7) which is a key regulator in retinol metabolism. Their results were consistent with our finding that RARs were downregulated with neonatal opioid exposure, and morphine exposure likely disturbed retinol metabolism in the gut therefore reduced transcription activities by RAR and RAR/RXR. The present study showed that the probiotics supplementation could upregulate the RAR and RXR that were downregulated by neonatal opioid exposure. Multiple studies in the field have demonstrated the direct involvement of gut microbiota in vitamin A metabolism. Grizotte-Lake et al^47^ demonstrated a direct role of commensal bacteria in modulating the concentration of the vitamin A metabolite retinoid acid in the gut. They reported that bacteria belonging to class Clostridia suppressed Rdh7 expression and retinoid acid synthesis. Woo et al^48^ found that segmented filamentous bacteria (SFB) and other beneficial commensal bacteria generate intestinal retinoic acid levels despite inhibition of host production. Bonakdar et al^49^ discovered that gut bacteria *Lactobacillus intestinalis* metabolized vitamin A and specifically restored retinoid acid levels in the gut of vancomycin-treated mice. Our previous research had shown that NME adolescents given *B. infantis* supplementation showed an increase in mainly the commensal genera of *Lactobacillus*, *Bacteroides,* and *Turicibacter*, which may play a role in regulating vitamin A metabolism and RAR/RXR activities ^4^.

Although this study provides important insights into the molecular and cellular consequences of NME, it has several limitations that warrant consideration. While the use of a murine model allowed for rigorous experimental control and mechanistic insight into the effects of NME and microbiome-targeted interventions, the translational relevance of these findings remains to be established. Future clinical studies are needed to assess whether probiotic supplementation can similarly mitigate opioid-induced neurodevelopmental disruptions in human infants. Moreover, while this study focused primarily on pain-related pathways, early-life opioid exposure has been associated with a broader range of neurobehavioral alterations. Follow-up studies examining additional behavioral domains, such as locomotor activity, learning, and cognition, will be critical to fully elucidate the long-term functional consequences of NME. Lastly, our use of the 10x Genomics Chromium Fixed RNA Profiling assay, which is a probe-based single-cell transcriptomic platform, limited our ability to detect non-coding RNAs and novel transcripts not included in the predefined probe set. Future work using whole-transcriptome approaches may uncover additional regulatory mechanisms underlying the observed effects.

In summary, this study marks a significant advance as the first to present single-cell RNA sequencing data from the adolescent midbrain following neonatal morphine exposure and probiotic intervention. Using a murine model, we demonstrated that neonatal morphine exposure profoundly disrupted cell-type-specific gene expression, canonical signaling pathways, and intercellular communication within the adolescent midbrain. Notably, probiotic supplementation with *B. infantis* effectively reversed many of these morphine-induced alterations, including the downregulation of Hox genes across multiple cell types and the activation of neuropathic pain signaling pathways. These findings offer unprecedented molecular and cellular insights into the long-term neurodevelopmental effects of early opioid exposure. Furthermore, they highlight the therapeutic potential of microbiome-targeted interventions as a promising strategy to counteract opioid-induced molecular reprogramming and promote neurodevelopmental resilience—opening new avenues for treating early-life opioid-related deficits.

## CRediT authorship contribution statement

**Junyi Tao:** Conceptualization, Software, Formal analysis, Visualization, Data curation, Writing – original draft. **Danielle Antoine**: Conceptualization, Writing – review & editing, Visualization, Methodology, Investigation, Resources. **Richa Jalodia:** Resources, Investigation. **Eridania Valdes:** Resources, Investigation. **Sean Michael Boyles**: Resources, Investigation. **William Hulme**: Resources, Investigation. **Sabita Roy**: Conceptualization, Writing – review & editing, Supervision, Funding acquisition, Conceptualization.

## Data Availability

The raw single cell RNA sequencing data has been deposited to NCBI SRA with accession number PRJNA1260540 https://dataview.ncbi.nlm.nih.gov/object/PRJNA1260540?reviewer=u2bc5vm45kvcavo56i3piem l0v.

## Acknowledgment

We would like to thank Nicolas Alberti (University of Miami) for helping to set up the supercomputer (Pegasus Cluster) and installing packages and dependencies. We would like to that Institute for Data Science & Computing at University of Miami for the service units provided by Early Career Researcher Award (EC202303).

