## Supplemental Table S1-S7 for "Single-cell transcriptomics reveals probiotic reversal of neonatal morphine-induced gene disruptions underlying adolescent pain hypersensitivity"

**Supplementary materials**

**Supplemental Figures**

**Figure S1. Cellular composition by group and by sample, related to Figure 1**. (A). Barplot showing cellular composition by treatment group and sample. (B). UMAP plot labelled by treatment groups after integration. (C). UMAP plot labelled by sample after integration. (D). UMAP plot labelled by cell types and split by conditions.

**Figure S2. Side-by-side DEGs comparisons** of Oligodendrocyte precursor cells (A), microglia cells (B), and Cholinergic neurons (C). in Mor vs Sal, Mor+Pro vs Mor and Mor+Pro vs Sal conditions. Genes with $|Log2FC| > 0.25$ and adjusted p value $< 0.05$ are counted as significantly changed. Related to Figure 4. M vs S, morphine vs saline. MP+M, morphine +probiotics vs morphine. MP vs S, morphine+probiotics vs saline.

**Figure S3. Overall neuropathic pain pathways** of Oligodendrocytes under Mor vs Sal (A) and Mor+Pro vs Mor (B) conditions. Related to Figure 6. M vs S, morphine vs saline. MP+M, morphine +probiotics vs morphine.

**Figure S4. Detailed views of glutamate transporters subunits in neuropathic pain pathways** of Glutamatergic neurons under Mor vs Sal (A) and Mor+Pro vs Mor (B) conditions. Astrocytes under Mor vs Sal (C) and Mor+Pro vs Mor (D) conditions. Oligodendrocytes under Mor vs Sal (E) and Mor+Pro vs Mor (F) conditions. Related to Figure 6. M vs S, morphine vs saline. MP+M, morphine +probiotics vs morphine.

**Figure S5. cell–cell communication among Glutamatergic neurons, astrocytes and oligodendrocytes. related to Figure 7.** (A) Circle plot showing differential interaction strength between Glutamatergic neurons, astrocytes and oligodendrocytes under M vs S. Red (or blue) colored edges represent increased (or decreased) signaling in the comparison. (B) Circle plot showing differential interaction strength between Glutamatergic neurons, astrocytes and oligodendrocytes under MP+M. (C) Overall information flow of top signaling pathway from astrocytes and oligodendrocytes to glutamatergic neuron under M vs S. (D) Overall information flow of top signaling pathway from astrocytes and oligodendrocytes to glutamatergic neuron under MP+M. M vs S, morphine vs saline. MP+M, morphine +probiotics vs morphine.

**Figure S6. IPA Upstream Regulator Analysis** of Glutamatergic neurons under Mor vs Sal (A) and Mor+Pro vs Mor (B) conditions, GABAergic neurons under Mor vs Sal (C) and Mor+Pro vs Mor (D) conditions, related to Figure 8. M vs S, morphine vs saline. MP+M, morphine +probiotics vs morphine.

**Figure S7. IPA Upstream Regulator Analysis** of Astrocytes under Mor+Pro vs Mor conditions, related to Figure 8. MP+M, morphine +probiotics vs morphine.

**Supplemental Tables**

**Table S1. DEGs of all cell types under Mor vs Sal and Mor+Pro vs Mor.**

**Table S2. IPA canonical pathway results for glutamatergic neurons, GABAergic neurons, astrocytes and oligodendrocytes under Mor vs Sal and Mor+Pro vs Mor.**

**Table S3. IPA upstream analysis results for glutamatergic neurons, GABAergic neurons, astrocytes and oligodendrocytes under Mor vs Sal and Mor+Pro vs Mor.**

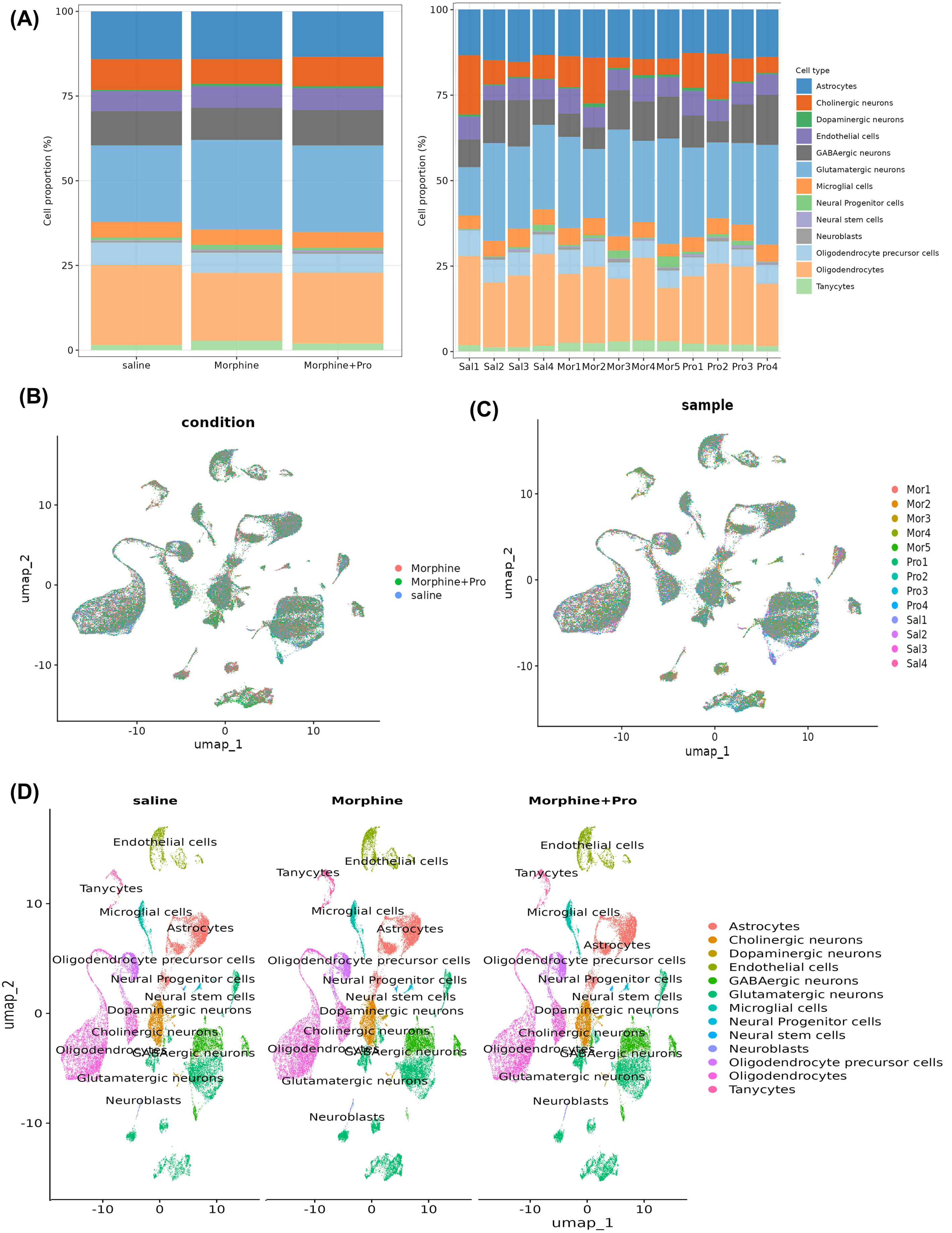

(A)
Cell proportion (%)
100
75
50
25
0
saline
Morphine
Morphine+Pro
Sal1
Sal2
Sal3
Sal4
Mor1
Mor2
Mor3
Mor4
Mor5
Pro1
Pro2
Pro3
Pro4
Cell type
Astrocytes
Cholinergic neurons
Dopaminergic neurons
Endothelial cells
GABAergic neurons
Glutamatergic neurons
Microglial cells
Neural Progenitor cells
Neural stem cells
Neuroblasts
Oligodendrocyte precursor cells
Oligodendrocytes
Tanycytes
(B)
condition
umap_2
umap_1
10
0
-10
Morphine
Morphine+Pro
saline
(C)
sample
umap_2
umap_1
Mor1
Mor2
Mor3
Mor4
Mor5
Pro1
Pro2
Pro3
Pro4
Sal1
Sal2
Sal3
Sal4
(D)
saline
Morphine
Morphine+Pro
umap_2
umap_1
Endothelial cells
Tanycytes
Microglial cells
Astrocytes
Oligodendrocyte precursor cells
Neural Progenitor cells
Neural stem cells
Dopaminergic neurons
Cholinergic neurons
Oligodendrocytes
GABAergic neurons
Glutamatergic neurons
Neuroblasts
Astrocytes
Cholinergic neurons
Dopaminergic neurons
Endothelial cells
GABAergic neurons
Glutamatergic neurons
Microglial cells
Neural Progenitor cells
Neural stem cells
Neuroblasts
Oligodendrocyte precursor cells
Oligodendrocytes
Tanycytes


**Figure S1**

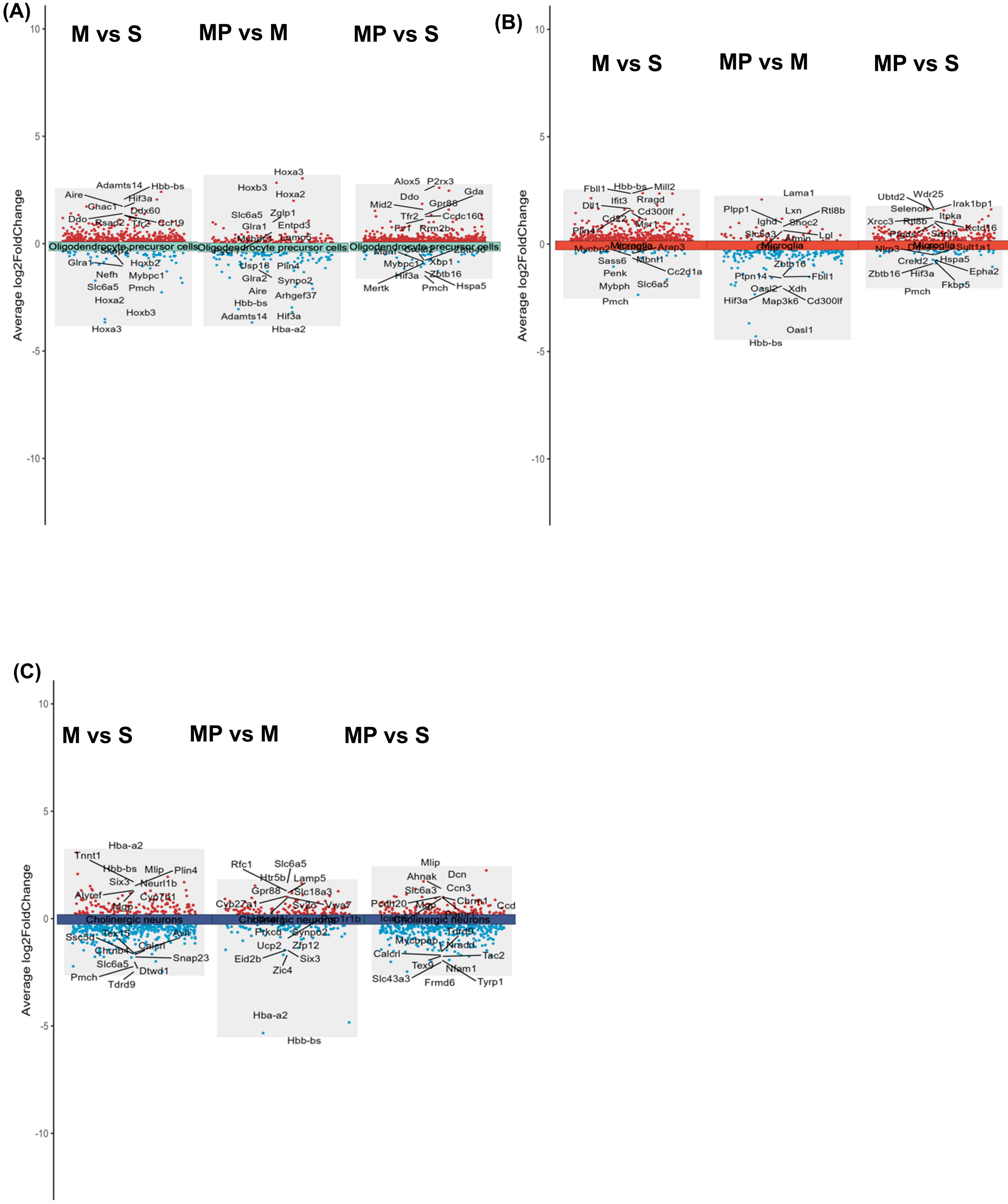
(A)
M vs S
MP vs M
MP vs S
Average log2FoldChange
10
5
0
-5
-10
Oligodendrocyte precursor cells
Adamts14
Hbb-bs
Aire
Hif3a
Ghac1
Ddx60
Ddo
Rsad2
Tfr2
Ccl19
Glra1
Hoxb2
Nefh
Mybpc1
Slc6a5
Pmch
Hoxa2
Hoxb3
Hoxa3
Hoxa3
Hoxb3
Hoxa2
Slc6a5
Zglp1
Glra1
Entpd3
Lamp5
Usp18
Plin4
Glra2
Synpo2
Aire
Arhgef37
Hbb-bs
Adamts14
Hif3a
Hba-a2
Alox5
P2rx3
Gda
Ddo
Gpr88
Mid2
Tfr2
Ccdc160
Rrm2b
Xbp1
Mybpc1
Hif3a
Zbtb16
Mertk
Pmch
Hspa5
(B)
M vs S
MP vs M
MP vs S
Microglia
Fbll1
Hbb-bs
Mill2
Ifit3
Rragd
Dll1
Cd300lf
Cd22
Msr1
Plin4
Mycbp2
Arap3
Sass6
Mbnl1
Penk
Cc2d1a
Mybph
Slc6a5
Pmch
Lama1
Plpp1
Lxn
Rtl8b
Ighd
Shoc2
Slc6a3
Lpl
Zbtb16
Ptpn14
Fbll1
Oasl2
Xdh
Hif3a
Map3k6
Cd300lf
Oasl1
Hbb-bs
Ubtd2
Wdr25
Irak1bp1
Selenoh
Itpka
Xrcc3
Rtl8b
Kctd16
Nlrp3
Sult1a1
Creld2
Hspa5
Zbtb16
Hif3a
Epha2
Fkbp5
Pmch
(C)
M vs S
MP vs M
MP vs S
Cholinergic neurons
Hba-a2
Tnnt1
Hbb-bs
Mlip
Plin4
Six3
Neurl1b
Alyref
Cyp7b1
Mgp
Ssc5d
Tex15
Calcrl
Snap23
Slc6a5
Dtwd1
Pmch
Tdrd9
Rfc1
Slc6a5
Htr5b
Lamp5
Gpr88
Slc18a3
Cyp27a1
Sv2c
Vwa7
Prkcd
Synpo2
Ucp2
Zfp12
Eid2b
Six3
Zic4
Hba-a2
Hbb-bs
Mlip
Ahnak
Dcn
Slc6a3
Ccn3
Pcdh20
Mgp
Chrm1
Ccd
Tdrd9
Mycbpap
Nradd
Calcrl
Tac2
Tex9
Nfam1
Slc43a3
Frmd6
Tyrp1

**Figure S2**

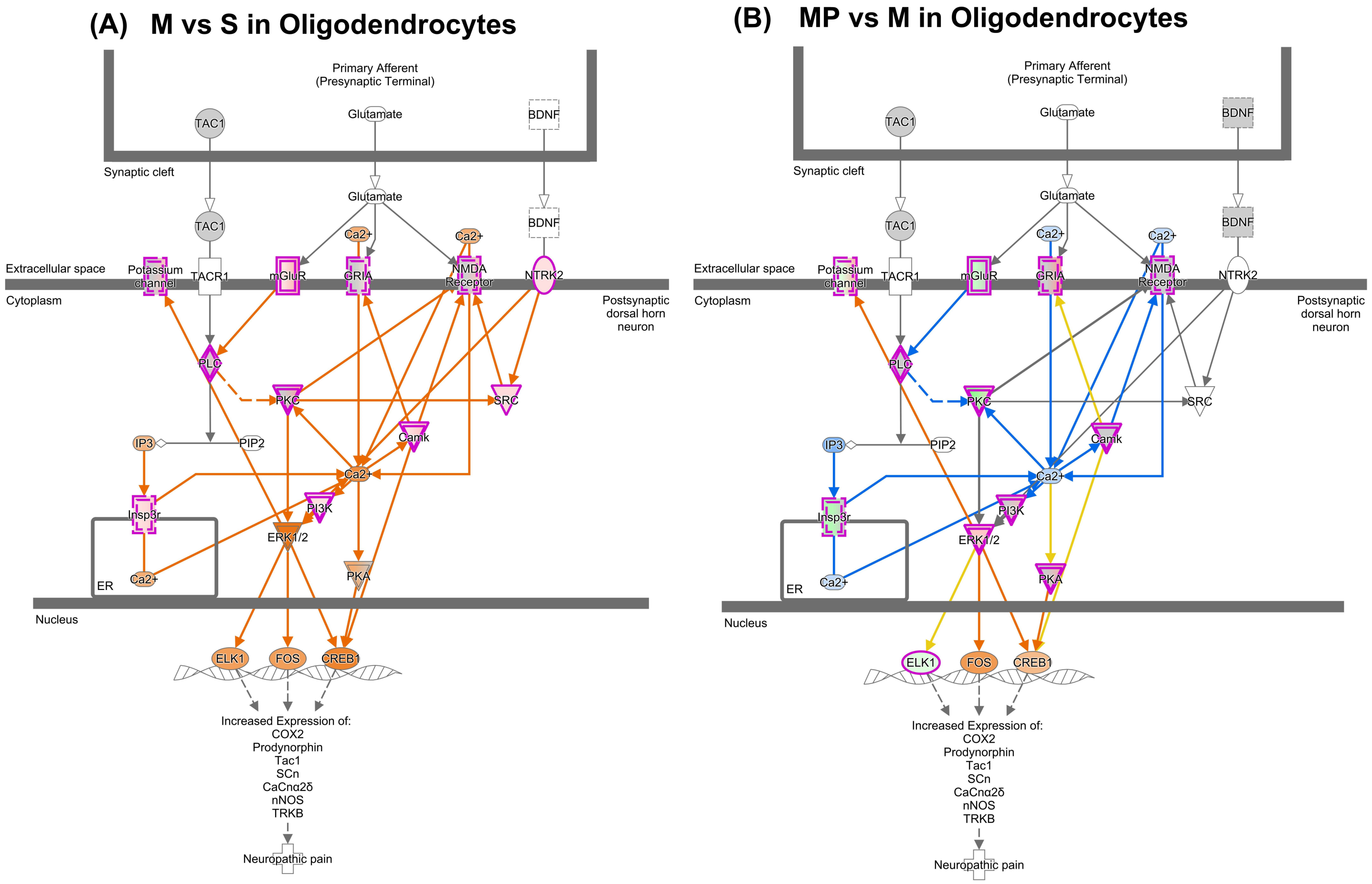
(A) M vs S in Oligodendrocytes
(B) MP vs M in Oligodendrocytes
Primary Afferent
(Presynaptic Terminal)
TAC1
Glutamate
BDNF
Synaptic cleft
Ca2+
Extracellular space
Cytoplasm
Potassium channel
TACR1
mGluR
GRIA
NMDA Receptor
NTRK2
Postsynaptic dorsal horn neuron
PLC
PKC
SRC
IP3
PIP2
Camk
Insp3r
PI3K
ERK1/2
PKA
ER
Nucleus
ELK1
FOS
CREB1
Increased Expression of:
COX2
Prodynorphin
Tac1
SCn
CaCnα2δ
nNOS
TRKB
Neuropathic pain

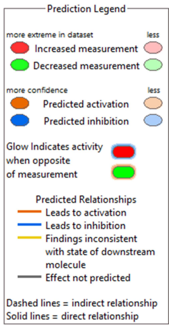
Prediction Legend
more extreme in dataset
less
Increased measurement
Decreased measurement
more confidence
less
Predicted activation
Predicted inhibition
Glow indicates activity when opposite of measurement
Predicted Relationships
Leads to activation
Leads to inhibition
Findings inconsistent with state of downstream molecule
Effect not predicted
Dashed lines = indirect relationship
Solid lines = direct relationship

Figure S3

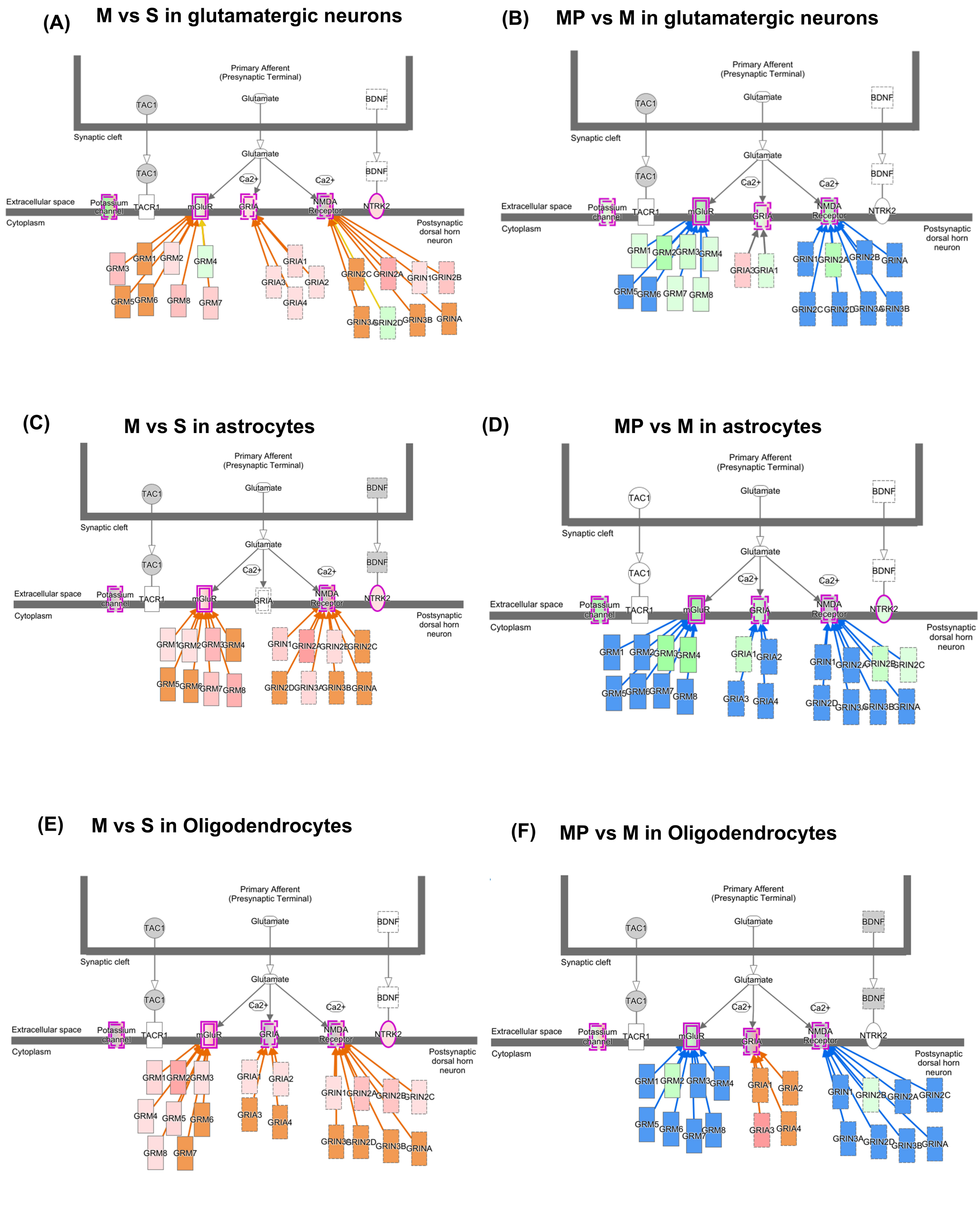
(A) M vs S in glutamatergic neurons
Primary Afferent (Presynaptic Terminal)
TAC1
Glutamate
BDNF
Synaptic cleft
Glutamate
TAC1
BDNF
Ca2+
Ca2+
Extracellular space
Potassium channel
TACR1
mGluR
GRIA
NMDA Receptor
NTRK2
Cytoplasm
Postsynaptic dorsal horn neuron
GRM1 GRM2 GRM3 GRM4 GRM5 GRM6 GRM7 GRM8
GRIA1 GRIA2 GRIA3 GRIA4
GRIN2C GRIN2A GRIN1 GRIN2B GRIN3A GRIN2D GRIN3B GRINA
(B) MP vs M in glutamatergic neurons
Primary Afferent (Presynaptic Terminal)
TAC1
Glutamate
BDNF
Synaptic cleft
Glutamate
TAC1
BDNF
Ca2+
Ca2+
Extracellular space
Potassium channel
TACR1
mGluR
GRIA
NMDA Receptor
NTRK2
Cytoplasm
Postsynaptic dorsal horn neuron
GRM1 GRM2 GRM3 GRM4 GRM5 GRM6 GRM7 GRM8
GRIA3 GRIA1
GRIN1 GRIN2A GRIN2B GRINA GRIN2C GRIN2D GRIN3A GRIN3B
(C) M vs S in astrocytes
Primary Afferent (Presynaptic Terminal)
TAC1
Glutamate
BDNF
Synaptic cleft
Glutamate
TAC1
BDNF
Ca2+
Ca2+
Extracellular space
Potassium channel
TACR1
mGluR
GRIA
NMDA Receptor
NTRK2
Cytoplasm
Postsynaptic dorsal horn neuron
GRM1 GRM2 GRM3 GRM4 GRM5 GRM6 GRM7 GRM8
GRIN1 GRIN2A GRIN2B GRIN2C GRIN2D GRIN3A GRIN3B GRINA
(D) MP vs M in astrocytes
Primary Afferent (Presynaptic Terminal)
TAC1
Glutamate
BDNF
Synaptic cleft
Glutamate
TAC1
BDNF
Ca2+
Ca2+
Extracellular space
Potassium channel
TACR1
mGluR
GRIA
NMDA Receptor
NTRK2
Cytoplasm
Postsynaptic dorsal horn neuron
GRM1 GRM2 GRM3 GRM4 GRM5 GRM6 GRM7 GRM8
GRIA1 GRIA2 GRIA3 GRIA4
GRIN1 GRIN2A GRIN2B GRIN2C GRIN2D GRIN3A GRIN3B GRINA
(E) M vs S in Oligodendrocytes
Primary Afferent (Presynaptic Terminal)
TAC1
Glutamate
BDNF
Synaptic cleft
Glutamate
TAC1
BDNF
Ca2+
Ca2+
Extracellular space
Potassium channel
TACR1
mGluR
GRIA
NMDA Receptor
NTRK2
Cytoplasm
Postsynaptic dorsal horn neuron
GRM1 GRM2 GRM3 GRM4 GRM5 GRM6 GRM8 GRM7
GRIA1 GRIA2 GRIA3 GRIA4
GRIN1 GRIN2A GRIN2B GRIN2C GRIN3A GRIN2D GRIN3B GRINA
(F) MP vs M in Oligodendrocytes
Primary Afferent (Presynaptic Terminal)
TAC1
Glutamate
BDNF
Synaptic cleft
Glutamate
TAC1
BDNF
Ca2+
Ca2+
Extracellular space
Potassium channel
TACR1
mGluR
GRIA
NMDA Receptor
NTRK2
Cytoplasm
Postsynaptic dorsal horn neuron
GRM1 GRM2 GRM3 GRM4 GRM5 GRM6 GRM7 GRM8
GRIA1 GRIA2 GRIA3 GRIA4
GRIN1 GRIN2B GRIN2A GRIN2C GRIN3A GRIN2D GRIN3B GRINA

**Figure S4**

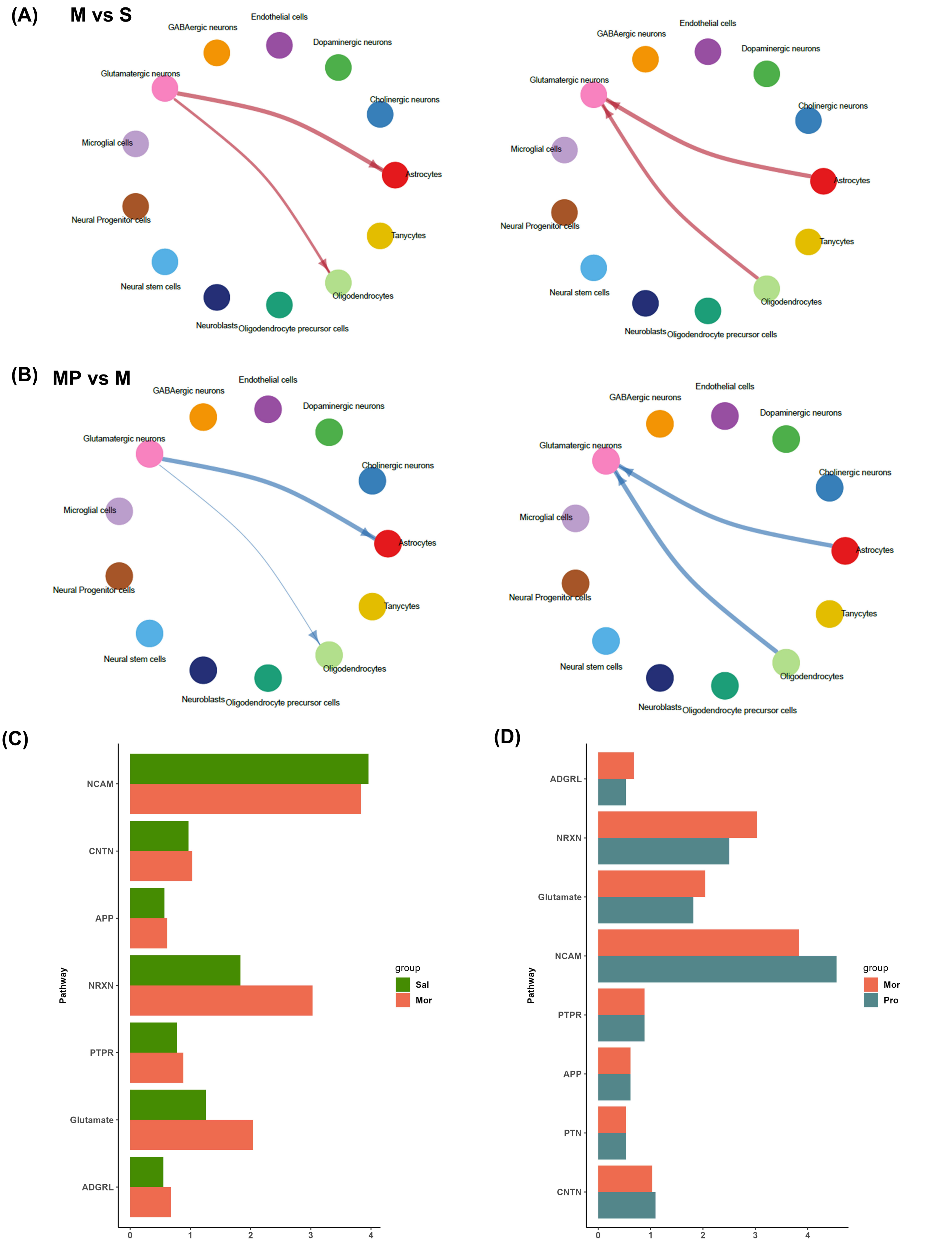

(A)
M vs S
GABAergic neurons
Endothelial cells
Dopaminergic neurons
Glutamatergic neurons
Cholinergic neurons
Microglial cells
Astrocytes
Neural Progenitor cells
Tanycytes
Neural stem cells
Oligodendrocytes
Neuroblasts
Oligodendrocyte precursor cells
(B)
MP vs M
(C)
NCAM
CNTN
APP
NRXN
PTPR
Glutamate
ADGRL
Pathway
Information flow
group
Sal
Mor
(D)
ADGRL
NRXN
Glutamate
NCAM
PTPR
APP
PTN
CNTN
Pathway
Information flow
group
Mor
Pro


Figure S5

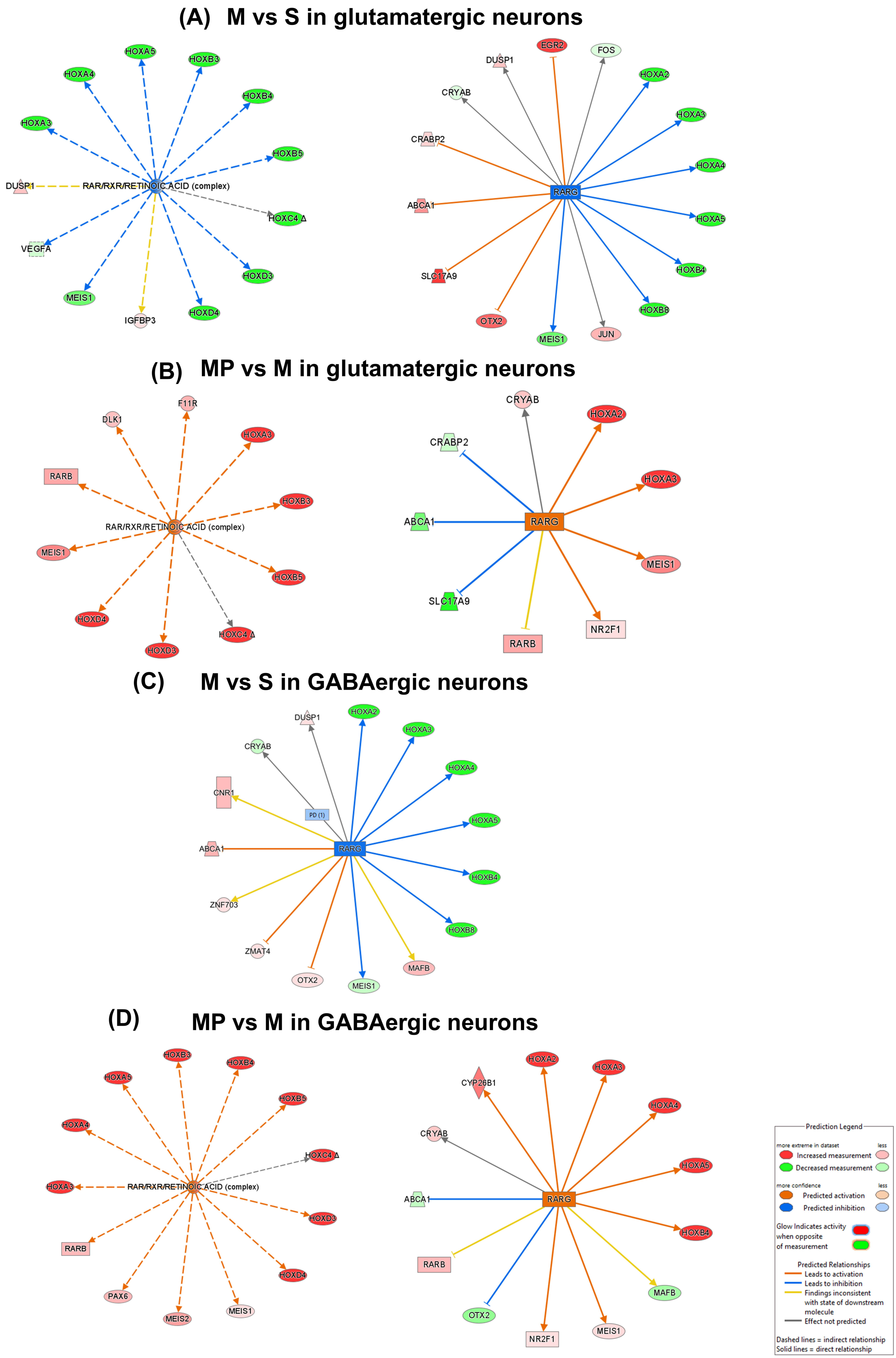
(A) M vs S in glutamatergic neurons
HOXA5
HOXB3
HOXA4
HOXB4
HOXA3
HOXB5
DUSP1
RAR/RXR/RETINOIC ACID (complex)
HOXC4 Δ
VEGFA
HOXD3
MEIS1
HOXD4
IGFBP3
EGR2
FOS
DUSP1
HOXA2
CRYAB
HOXA3
CRABP2
HOXA4
RARG
ABCA1
HOXA5
HOXB4
SLC17A9
HOXB8
OTX2
MEIS1
JUN
(B) MP vs M in glutamatergic neurons
F11R
DLK1
HOXA3
RARB
HOXB3
RAR/RXR/RETINOIC ACID (complex)
MEIS1
HOXB5
HOXD4
HOXC4 Δ
HOXD3
CRYAB
HOXA2
CRABP2
HOXA3
ABCA1
RARG
MEIS1
SLC17A9
NR2F1
RARB
(C) M vs S in GABAergic neurons
DUSP1
HOXA2
HOXA3
CRYAB
HOXA4
CNR1
PD (1)
HOXA5
ABCA1
RARG
HOXB4
ZNF703
HOXB8
ZMAT4
MAFB
OTX2
MEIS1
(D) MP vs M in GABAergic neurons
HOXB3
HOXB4
HOXA5
HOXB5
HOXA4
HOXC4 Δ
HOXA3
RAR/RXR/RETINOIC ACID (complex)
HOXD3
RARB
HOXD4
PAX6
MEIS2
MEIS1
HOXA2
CYP26B1
HOXA3
HOXA4
CRYAB
HOXA5
ABCA1
RARG
HOXB4
RARB
MAFB
OTX2
NR2F1
MEIS1
Prediction Legend
more extreme in dataset
less
Increased measurement
Decreased measurement
more confidence
less
Predicted activation
Predicted inhibition
Glow Indicates activity when opposite of measurement
Predicted Relationships
Leads to activation
Leads to inhibition
Findings inconsistent with state of downstream molecule
Effect not predicted
Dashed lines = indirect relationship
Solid lines = direct relationship

**Figure S6**

### MP vs M in astrocytes

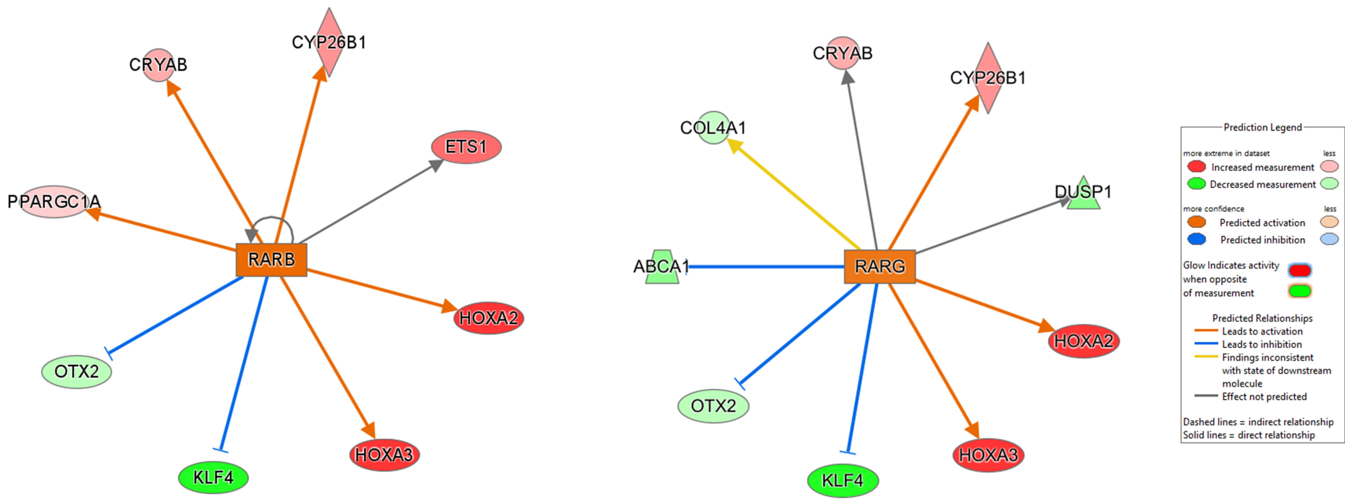


**Figure S7**
